# Characterization of full-length and cytoplasmic tail-truncated envelope glycoproteins incorporated into human immunodeficiency virus (HIV-1) virions and virus-like particles

**DOI:** 10.1101/2025.07.10.664145

**Authors:** Saumya Anang, Shijian Zhang, Amanda Ennis, Haitao Ding, Ashlesha Deshpande, Hanh T. Nguyen, John C. Kappes, Joseph G. Sodroski

## Abstract

During transport to the surface of infected cells, the human immunodeficiency virus (HIV-1) envelope glycoprotein (Env) trimer is cleaved to produce the mature functional Env trimer ((gp120/gp41)_3_). Env cleavage stabilizes the pretriggered Env conformation (PTC), the major target for broadly neutralizing antibodies. Although the mature Env is relatively enriched in virions and virus-like particles (VLPs), conformationally flexible uncleaved Envs typically contaminate preparations of these particles. In non-permissive cells, the long ∼149-residue gp41 cytoplasmic tail (CT) is necessary for Env incorporation into virions. In a minority of HIV-1 strains, the gp41 CT is clipped in virions by the viral protease. Here, we compare Envs with CT truncations and CT alterations that increase or decrease protease clipping in permissive cells. Changes in protease clipping affected amphotericin B sensitivity, but did not alter other viral phenotypes. By contrast, a corresponding CT truncation (L748STOP) increased cell-surface and virion Env levels, cell-cell fusion, and virus infectivity and cytotoxicity. Notably, in diverse HIV-1 strains, the ratio of cleaved/uncleaved Envs in preparations of virions and extracellular vesicles was increased by this CT truncation. ESCRT and ALIX-binding region (EABR) vesicles incorporated significantly more uncleaved CT-truncated Env than HIV-1 VLPs. Env CT deletion/truncation did not qualitatively alter the viral neutralization profile; however, increased antibody concentrations were required to neutralize viruses with the higher levels of cleaved Env that resulted from CT truncation. Specific CT truncations provide a means of enriching the PTC and limiting the incorporation of nonfunctional and conformationally heterogeneous uncleaved Envs into preparations of virions and VLPs.

**IMPORTANCE:** The human immunodeficiency virus (HIV-1) envelope glycoprotein (Env) trimer mediates entry of the virus into host cells. The pretriggered conformation (PTC) of Env is the major target for protective broadly neutralizing antibodies, but the PTC is unstable and therefore difficult to study. The cleavage of the flexible Env precursor stabilizes the PTC. Therefore, the presence of uncleaved Env compromises the purity of the PTC in Env preparations. We found that certain truncations of the Env cytoplasmic tail resulted in improved ratios of cleaved:uncleaved Env in preparations of HIV-1 viruses or virus-like particles. In some contexts, cytoplasmic tail truncation increased the level of Env in virus preparations. Although higher concentrations of antibodies were required to neutralize these viruses, Envs with specific truncations of the cytoplasmic tail retained the PTC. Thus, cytoplasmic tail truncation could assist efforts to purify and characterize the Env PTC on the viral membrane.

## INTRODUCTION

The human immunodeficiency virus (HIV-1) envelope glycoprotein (Env) trimer ((gp120/gp41)_3_) mediates virus entry and cytopathic effects (1–5). In the infected cell, a fraction of the Env precursor in the endoplasmic reticulum passes through the Golgi, where it is cleaved and modified by complex glycans on the way to the cell surface and virions (6–12). Prior to receptor binding, virion Envs largely occupy the pretriggered conformation (PTC), which is a major target for small-molecule entry inhibitors and broadly neutralizing antibodies (bNAbs) (13–16). Uncleaved Env, on the other hand, is conformationally flexible, recognized by poorly neutralizing antibodies (pNAbs) and antigenically distinct from the PTC (17–22). Therefore, uncleaved Env is an undesirable component in preparations used to study the structure and immunogenicity of the pretriggered Env.

The Envs of HIV-1 and other lentiviruses have unusually long cytoplasmic tails (CTs) (12,23,24). The ∼149-residue HIV-1 Env CT consists of a relatively unstructured CT_N_ region (residues 707-752) and a CT_C_ region (residues 753-856) that contains membrane-interactive amphipathic helices (12,23–26). The HIV-1 Env CT is palmitoylated and has endocytic motifs (^712^YXXL^715^ and a C-terminal dileucine motif [^855^LL^856^]), which interact with clathrin adaptor proteins (27–37). In non-permissive cells, which include primary target cells, the CT is important for the recruitment of Env to virus assembly sites and Env incorporation into HIV-1 virions (38–46). This process has been reported to involve the interaction of the Env CT with Rab 11 family-interacting protein 1C (FIP1C), a member of a family of proteins that mediate sorting of cargo from the endosomal recycling compartment to the plasma membrane (47–49). However, a more recent study used cells in which FIP1C expression was knocked out to show that virion incorporation in primary CD4^+^ T cells and in some cell lines is not dependent on FIP1C (50). In permissive cells, the CT is not required for Env incorporation into virions (38–44,46). Compatibility between the HIV-1 matrix (MA) protein and the Env CT can influence the efficiency of Env incorporation into virions (51,52). For example, some MA changes that exclude Env from virus particles can be compensated in permissive cells by CT truncation (53,54). Thus, both cell- and virus-specific factors influence the phenotypes of Env CT variants with respect to virion Env incorporation.

Alterations of the Env CT have been reported to affect HIV-1 sensitivity to neutralizing antibodies, host cell factors and antiviral agents. For some HIV-1 strains, CT truncation has been reported to alter Env antigenicity and decrease virus sensitivity to broadly neutralizing antibodies directed against quaternary V2 gp120 epitopes near the Env trimer apex (55,56). In other cases, however, the absence of the CT exerted little or no influence on susceptibility to neutralization by antibodies (17). Truncation of the CT has been reported to alter the immunogenicity of HIV-1 Env coexpressed with Gag in an mRNA vaccine (57). CT deletions have also been reported to allow HIV-1 to escape from the host restriction factors SERINC5 and IFITMs (58). In the virions of some HIV-1 strains, clipping of the gp41 CT by the viral protease occurs; CT clipping, as well as truncation of the CT, leads to resistance to the cholesterol-binding fungal antibiotic amphotericin B (59–62). Finally, certain truncations of the CT have been suggested to increase the triggerability of HIV-1 Env, rendering Env more resistant to T20 peptides that recognize gp41 intermediates in the virus entry process (63–65). The mechanisms underlying the observed effects of Env CT changes on HIV-1 sensitivity to inhibition are not well understood.

Here, we compare the phenotypes of HIV-1 Env mutants with two types of CT alterations: 1) CT changes that increase or decrease CT clipping by the viral protease; and 2) Introduction of stop codons in *env* that truncate the CT near the membrane or at the viral protease clip site (between residues 747-748) (60,62). Except for altered sensitivity to amphotericin B, CT changes at the viral protease clip site did not significantly alter the viral phenotypes. By contrast, CT-truncated Envs differed from the corresponding full-length Envs in several phenotypes determined in permissive virus-producing cells. Env variants with CT truncations were efficiently expressed on the cell surface and exhibited increases in the ratio of cleaved:uncleaved Env in preparations of virions, virus-like particles (VLPs) and extracellular vesicles (EVs). Viruses with these CT-truncated Envs exhibited higher levels of infectivity and cytopathic effects than viruses with full-length Envs. The antigenicity and glycosylation of the CT-truncated Envs were comparable to those of the full-length Envs, consistent with the qualitatively similar neutralization profiles of viruses with these Envs. Correlating with the increased levels of cleaved Env on the virions of some CT-truncated Env mutants, higher concentrations of entry blockers and antibodies were required to achieve neutralization. Relative to viruses with full-length Envs, the CT-truncated viruses were resistant to amphotericin B but were equally sensitive to methyl-β-cyclodextrin, BMS-806, SERINC5 and IFITM. These results suggest that specific CT truncations can be a useful means to increase the level of cleaved pretriggered HIV-1 Env in virion/VLP preparations.

## RESULTS

### HIV-1 Env cytoplasmic tail (CT) constructs and their processing

In this study, we compared the phenotypes of the full-length HIV-1_AD8_ Env with those of two types of Env CT mutants: 1) mutants with alterations in the CT site of viral protease clipping (between residues 747-748); and 2) CT-truncated Envs resulting from the introduction of stop codons in *env* (Fig. 1A). The Env of HIV-1_AD8_, a primary Tier-2 strain, is efficiently cleaved and therefore was included in most of these studies (11,17,21). The R747L and L748A changes increase and decrease, respectively, the clipping of the CT by the viral protease (60,62). The R747L and L748A changes should manifest their phenotypes primarily after Env has been incorporated into virions. The CT-truncated Envs were produced by introducing stop codons in place of the codons for Tyr 712, Leu 715 and Leu 748. The L748STOP Env has a CT that is truncated at a position corresponding to the site at which clipping of the full-length HIV-1_AD8_ Env by the viral protease occurs (60). The phenotypic effects of the CT truncations could potentially manifest in the Env-expressing cells as well as in the virions. The designed set of HIV-1_AD8_ Env mutants allows a direct comparison of the phenotypic effects of CT clipping by the viral protease in virions and CT truncation.

**FIG 1.**
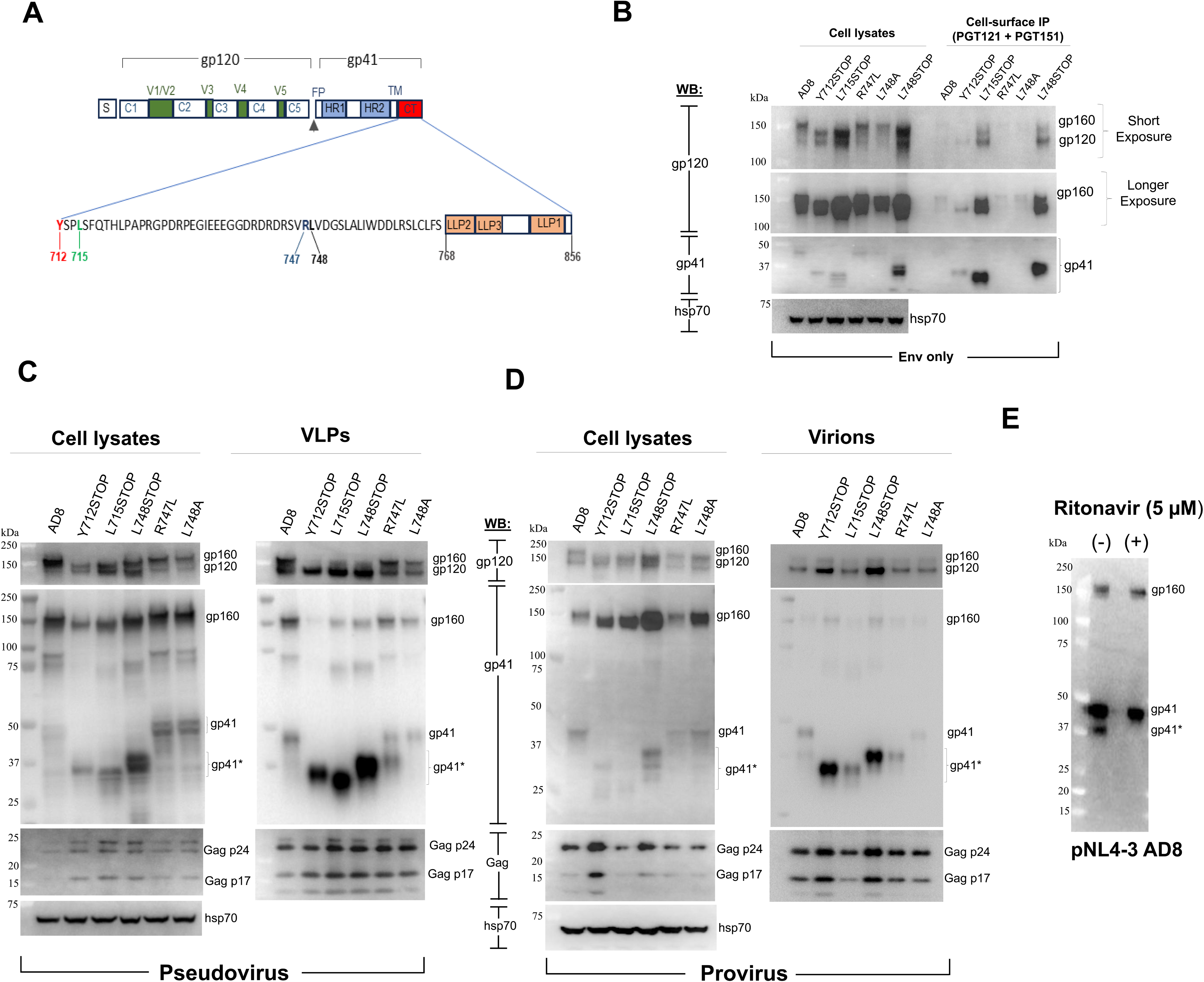
Expression and processing of HIV-1_AD8_ Env cytoplasmic tail (CT) mutants. (A) In the schematic representation of the HIV-1_AD8_ Env, the signal peptide (S) and the gp120 and gp41 subunits are shown. The gp120 constant (C1-C5) and variable (V1-V5) regions, gp120-gp41 cleavage site (black triangle), the gp41 fusion peptide (FP), heptad repeat (HR1 and HR2) regions and transmembrane (TM) region are depicted. The gp41 cytoplasmic tail (CT) sequence is shown in detail, with the lentivirus lytic peptide (LLP) regions depicted. The CT amino acid residues altered in this study are highlighted; the residues are numbered according to standard designation (131). (B) The expression of the wild-type HIV-1_AD8_ and mutant Env variants in total cell lysates and on cell surfaces was examined. HEK 293T cells were transfected with pSVIIIenv plasmids encoding the wild-type HIV-1_AD8_ Env or CT-altered variants along with a plasmid expressing the HIV-1 Tat protein in a DNA weight ratio of 8:1 (“Env only”). Forty-eight hours after transfection, cells were either lysed or used for cell-surface Env immunoprecipitation (IP) by incubation with 5 µg each of PGT121 and PGT151 antibodies for 1.5 hours at room temperature. The cells were then washed with 1x PBS and lysed, and the cell lysates were precipitated with Protein A-agarose beads for 1 hour at 37°C. The beads were washed three times and, along with the total cell lysates, Western blotted with a goat anti-gp120 antibody (first and second upper panels) and the 4E10 anti-gp41 antibody (third panel). The cell lysates were also Western blotted with a rabbit anti-hsp 70 antibody (bottom panel). (C,D) The expression and processing of the indicated Env variants in cell lysates and VLPs/virions were examined. In C, HEK 293T cells were transfected with pSVIIIenv plasmids expressing the wild-type HIV-1_AD8_ Env or CT-altered Env variants, an HIV-1 luciferase-expressing vector and the psPAX2 plasmid expressing HIV-1 packaging proteins (“Pseudovirus”). In D, HEK 293T cells were transfected with pNL4-3 proviral plasmids expressing the indicated HIV-1_AD8_ Env variants (“Provirus”). Forty-eight hours later, the cell supernatants were clarified by a low-speed spin, filtered through a 0.45-µm membrane, and centrifuged at 14,000 x g for 1 hour at 4°C. In parallel, the cells were lysed. The pelleted VLPs/virions and clarified cell lysates were Western blotted with a goat anti-gp120 antibody (upper panels), the 4E10 anti-gp41 antibody (middle panels) or antibodies against Gag p55/p24/p17 or hsp70 (lower panels). (E) To evaluate the contribution of the HIV-1 protease to clipping of the gp41 CT, the pNL4-3 AD8 proviral plasmid was transfected into HEK 293T cells, which were treated with 5 µM ritonavir. Viruses were harvested from cell supernatants 48 hours later and analyzed by Western blotting with the 4E10 anti-gp41 antibody. Transfected cells that were not treated with ritonavir were studied in parallel. In C-E, the multiple forms of gp41 resulting from clipping or truncation of the CT are labeled gp41*. The results shown are representative of those obtained in more than two independent experiments.

The Env variants with CT alterations and truncations were expressed with Rev (“Env only”), in recombinant HIV-1 pseudotypes (“Pseudovirus”) or with a full complement of other HIV-1 proteins in infectious molecular proviral clones (IMCs) (“Provirus”) (Fig. 1B-D). Env expression was evaluated by cotransfection of HEK 293T cells with the pSVIIIenv expressor plasmids and a plasmid encoding the HIV-1 Tat protein (8:1 Env:Tat weight ratio) (Fig. 1B). Forty-eight hours later, Envs in cell lysates were analyzed by Western blotting. Envs on the cell surface were precipitated with a mixture of the PGT121 and PGT151 antibodies. The PGT121 antibody recognizes a glycan-dependent gp120 epitope near the third variable (V3) loop, and the PGT151 antibody recognizes a glycan-dependent epitope spanning the gp120-gp41 interface (66,67). The CT-truncated L715STOP and L748STOP mutants were expressed at the highest levels in cell lysates and on the cell surface (Fig. 1B). The ratio of cleaved:uncleaved Env was higher for the CT-truncated Envs (Y712STOP, L715STOP and L748STOP) than for the wild-type AD8, R747L and L748A Envs.

We next evaluated the processing and incorporation of these Envs into pseudotyped VLPs. Pseudovirus VLPs were produced from HEK 293T cells cotransfected with pSVIIIenv, the psPAX2 packaging plasmid and a luciferase-expressing HIV-1 vector (Fig. 1C). Although all of the Env variants were efficiently expressed, the overall Env level in both the cell lysates and the VLPs was greatest for the L748STOP Env. Most of the Envs in the cell lysates were uncleaved, whereas the cleaved Envs were enriched in the VLPs. The ratio of cleaved:uncleaved Env in the VLPs was similar for the wild-type AD8, R747L and L748A Envs. By contrast, the CT-truncated Envs exhibited much higher ratios of cleaved:uncleaved Env in the VLPs (see Fig. 2B below for quantitation). Clipping of the gp41 CT in the VLPs was increased by the R747L change and decreased by the L748A change, relative to that of the wild-type AD8 Env, as expected (62). Thus, CT truncations can increase the processing and incorporation of Env into pseudovirus VLPs.

**FIG 2.**
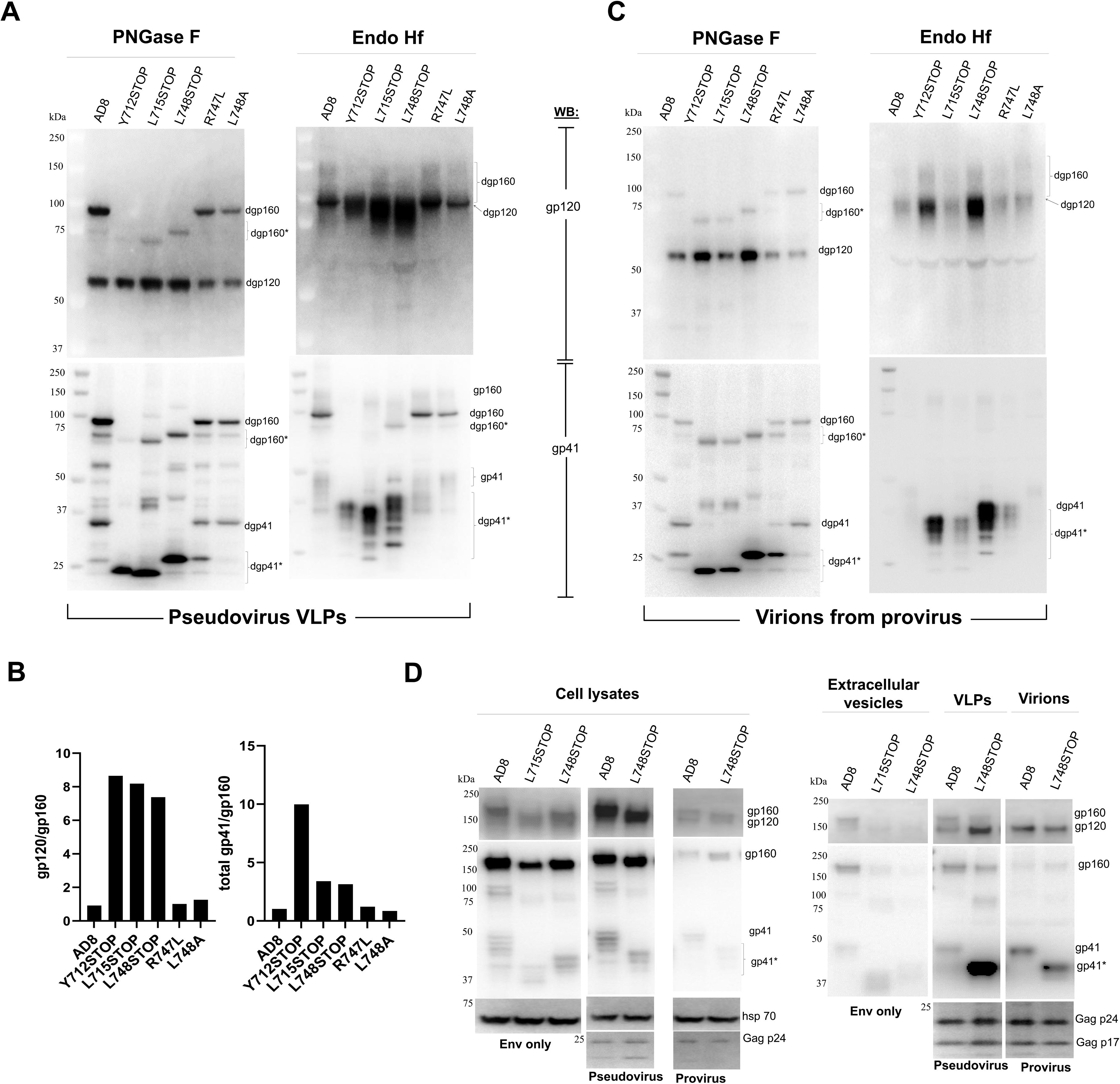
Glycosylation and processing of HIV-1_AD8_ and CT-altered Envs. (A) Pseudovirus VLPs produced in the experiment shown in Figure 1C were lysed and treated with PNGase F or Endo Hf. The deglycosylated Envs were analyzed by Western blotting with a goat anti-gp120 antibody (upper panels) and with the 4E10 anti-gp41 antibody (lower panels). Deglycosylated forms of Env are designated with a “d” (e.g., dgp160). CT-truncated or CT-clipped forms of gp160 and gp41 are asterisked. (B) Western blots of the PNGase F digests in A were used to measure the ratio of gp120/gp160 and the ratio of total gp41(clipped and unclipped)/gp160 on VLPs pseudotyped by the indicated Env variants. (C) Virions produced from transfected proviruses (IMCs) in the experiment shown in Figure 1D were lysed and treated with PNGase F or Endo Hf. The deglycosylated Envs were analyzed by Western blotting as described in A. Deglycosylated Env forms are designated with a “d” (e.g., dgp120). CT-truncated or CT-clipped forms of gp160 and gp41 are asterisked. The results shown in A, B and C are representative of those obtained in three independent experiments. (D) The expression and processing of full-length HIV-1_AD8_ and CT-truncated Envs in the lysates and supernatants of cells expressing Env only, pseudovirus VLPs, and virions produced from proviruses were compared. HEK 293T cells were transfected with the following: 1) pSVIIIenv plasmids expressing the indicated Env variants along with a Tat-expressing plasmid at an 8:1 DNA weight ratio (“Env only”); 2) pSVIIIenv Env-expressing plasmids and a psPAX2 plasmid expressing HIV-1 packaging proteins (“Pseudovirus”); or 3) pNL4-3 proviral plasmids expressing the indicated Envs (“Provirus”). Forty-eight hours later, the cell supernatants were clarified by low-speed centrifugation. The pseudovirus and provirus-produced virions were filtered through a 0.45-µm membrane. The cell supernatants were all centrifuged at 14,000 x g for 1 hour at 4°C. In parallel, the cells were lysed. Clarified cell lysates and the pellets from the cell supernatants (containing extracellular vesicles, VLPs and virions) were Western blotted as described in the Figure 1D legend. The results shown are representative of those obtained in three independent experiments.

Next, we examined the effects of CT changes on the quantity and quality of Env on virions produced by an infectious molecular proviral clone (IMC). The migration of the wild-type AD8, R747L and L748A gp41 glycoproteins in virions was consistent with partial viral protease clipping for the wild-type AD8 Env, nearly complete clipping for the R747L Env, and minimal clipping for the L748A Env (Fig. 1D). For all the Env variants, a higher ratio of cleaved:uncleaved Env was observed on virions than in cell lysates, consistent with previous observations that the processing of virion/VLP Envs produced from IMCs or defective proviruses is efficient (68–70). These results indicate that even in the case of virions produced by IMCs, which typically are enriched in mature Envs (68–70), CT-truncated Envs are efficiently expressed and maintain a high ratio of cleaved:uncleaved Env.

We confirmed the mechanism of gp41 CT clipping (60,62) by treating HEK 293T cells transfected with an IMC proviral plasmid expressing the wild-type HIV-1_AD8_ Env with ritonavir, an inhibitor of the HIV-1 protease (71). Cells transfected with the proviral plasmid that were not treated with ritonavir were studied in parallel. The virions from the untreated and ritonavir-treated cells were pelleted and lysed for Western blotting. The partial clipping of the gp41 glycoprotein observed for the wild-type HIV-1_AD8_ Env was not observed after ritonavir treatment (Fig. 1E). These results are consistent with the clipping of the HIV-1_AD8_ gp41 CT in virions being mediated by the viral protease (60,62).

### Characterization of Env CT mutants on VLPs, virions and extracellular vesicles

We evaluated the glycosylation of the HIV-1_AD8_ Env CT variants in pseudovirus VLPs and virions produced from IMCs. We treated lysates of VLPs and virions with protein N-glycosidase F (PNGase F), which removes all N-linked glycans, or endoglycosidase Hf (Endo Hf), which removes high-mannose and hybrid glycans but not complex glycans (72), and then analyzed the samples by Western blotting (Fig. 2A-C). The PNGase F digests confirmed that the CT truncations increased Env levels on VLPs/virions and also improved the ratio of cleaved:uncleaved Env on pseudovirus VLPs. The glycosylation patterns of the wild-type HIV-1_AD8_ Env and the CT variants were similar. The cleaved Envs on VLPs/virions were all Endo Hf-resistant, indicating that these Envs had passed though the Golgi compartment and had been modified by complex glycans. The small amount of gp160 on virions produced from IMCs was mostly Endo Hf-resistant and therefore modified by complex carbohydrates (Fig. 2C). Apparently, this represents gp160 glycoproteins that have passed through the Golgi compartment but escaped furin cleavage. Endo Hf-resistant gp160 was also observed on pseudovirus VLPs. In addition, the pseudovirus VLP preparations with the wild-type AD8, R747L and L748A Envs contained Endo Hf-sensitive gp160 that had not been modified by complex glycans (Fig. 2A, lower right panel). This high-mannose/hybrid glycan-containing gp160 may be a component of extracellular vesicles that contaminate the pseudovirus VLP preparations (see below). In summary, all of the cleaved and most of the uncleaved Envs on virions, regardless of Env CT structure, have acquired complex carbohydrates and therefore have passed through the Golgi network.

The above studies suggested that truncations of the Env CT could potentially be useful in preparing VLPs and virions with either increased Env levels or improved cleaved:uncleaved Env ratios. We compared the expression and processing of the wild-type AD8 and CT-truncated Envs in pseudovirus VLPs, provirus-produced virions and extracellular vesicles (EVs). EVs were prepared from the medium of cells expressing Env but no Gag proteins. The expression and processing of the full-length AD8 and CT-truncated Envs in the cell lysates for all systems were comparable (Fig. 2D, left panel). The EVs incorporated more wild-type AD8 Env than the L715STOP or L748STOP Envs (Fig. 2D, right panel). Most of the AD8 Env in EVs was uncleaved. Although the CT-truncated Envs were incorporated into EVs very inefficiently, these Envs were mostly cleaved in the EV preparations. In the pseudovirus VLPs, the L748STOP Env exhibited higher levels and a better cleaved:uncleaved Env ratio compared with the wild-type AD8 Env. Both the wild-type AD8 and L748STOP Envs in virions produced from a provirus were efficiently cleaved. Our results underscore the advantage of the L748STOP Env in producing pseudotyped VLP preparations with higher levels of mature Env from transiently expressing HEK 293T cells. The quality of the L748STOP pseudovirus VLPs may be helped by the negligible incorporation of CT-truncated Envs into EVs that potentially contaminate these VLP preparations.

### Incorporation of CT-truncated Envs and ESCRT- and ALIX-binding region (EABR)-tagged Envs into pseudovirus VLPs and EVs

Membrane proteins can be induced to self-assemble into EVs that bud from the cell membrane by directly recruiting proteins from the <u>E</u>ndosomal <u>S</u>orting <u>C</u>omplex <u>R</u>equired for <u>T</u>ransport (ESCRT) (73). This recruitment is mediated by a short peptide called the <u>E</u>SCRT- and <u>A</u>LIX-<u>b</u>inding <u>r</u>egion (EABR) positioned within the cytoplasmic tail of the membrane protein. We compared the quantity and quality of CT-truncated HIV-1_AD8_ Env variants incorporated into EABR EVs and pseudovirus VLPs. We added the EABR sequence to the HIV-1_AD8_ Env CT at positions 715 and 753 to allow comparison with the HIV-1_AD8_ L715STOP and L748STOP Envs, respectively. We also added the EABR sequence at CT position 753 of the TFAR Env, which is an HIV-1_AD8_ derivative stabilized in the PTC (69,70). These Env variants were expressed alone or together with the HIV-1 psPAX2 packaging plasmid and an HIV-1 luciferase-expressing vector in HEK 293T cells. Cell lysates were prepared, and cell supernatants were cleared by low-speed centrifugation and then centrifuged at 14,000 x g for 1 hour at 4°C. Lysates of cells and pellets prepared from the supernatants were analyzed by Western blotting, either directly (Fig. 3A) or after PNGase F/Endo Hf digestion (Fig. 3B).

**FIG 3.**
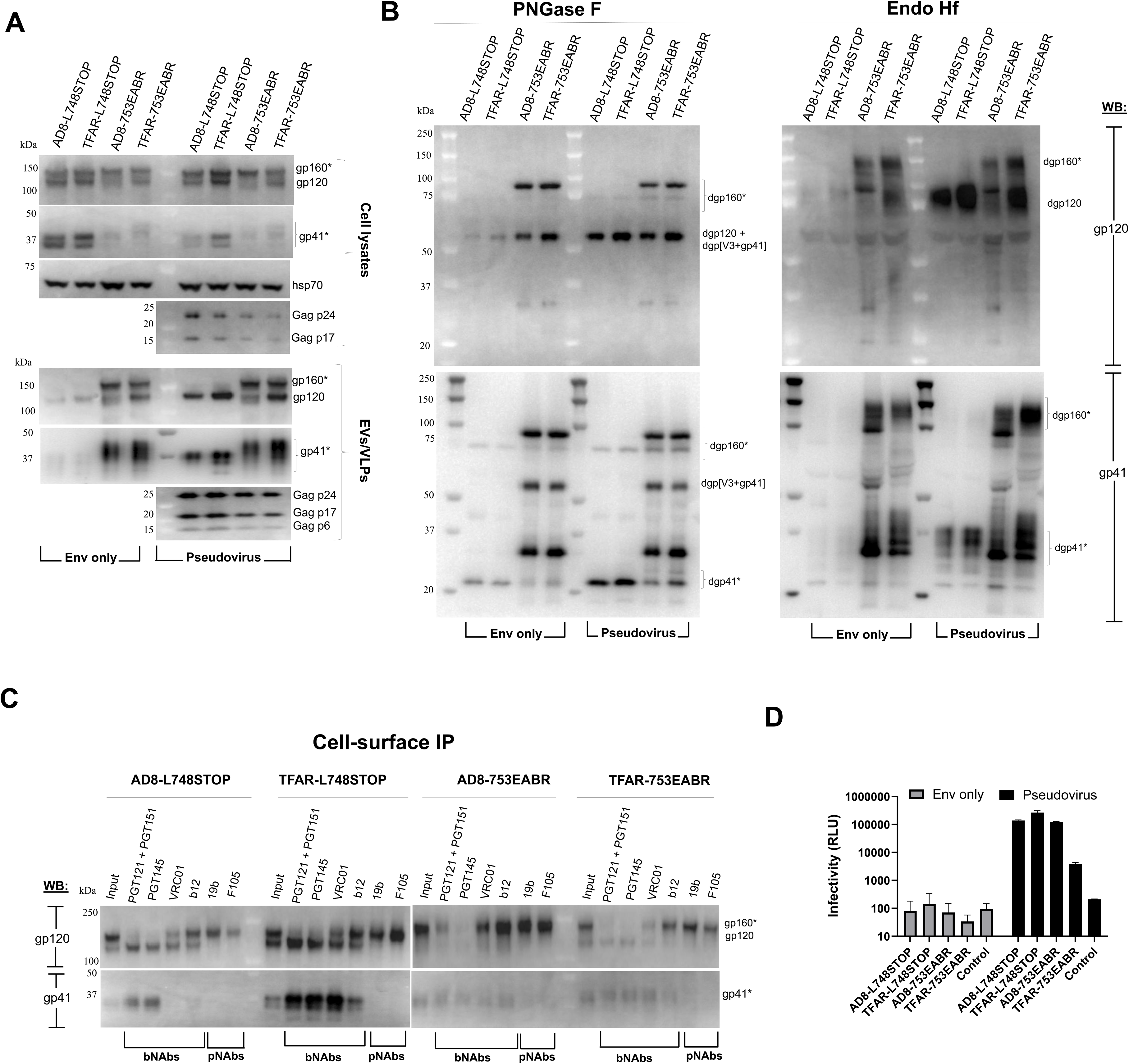
Comparison of CT-truncated Env and EABR-tagged Env incorporation into pseudovirus VLPs and EVs. HEK 293T cells were transfected with plasmids expressing the indicated HIV-1_AD8_ and TFAR Env variants, either alone (“Env only”) or with an HIV-1 luciferase-expressing vector and the psPAX2 packaging plasmid (“Pseudovirus”). Forty-eight hours later, the cells were lysed. Cell supernatants were clarified (600 x g for 10 minutes). The pseudovirus preparation was filtered (0.45-µm). EVs and pseudovirus VLPs were centrifuged at 14,000 x g for one hour at 4°C. Clarified cell lysates and lysed pellets from the cell supernatants (containing EVs and/or VLPs) were prepared. (A) The cell lysates and lysed EVs/VLPs were Western blotted as described in the Figure 1D legend. (B) The lysates of EVs from the supernatants of cells expressing only Env and the lysates of pseudovirus VLPs were treated with PNGase F or Endo Hf and Western blotted. The Western blots in the upper panels were developed with a goat anti-gp120 antibody and those in the lower panels with the 4E10 anti-gp41 antibody. The ∼54-kDa band (labeled dgp[V3+gp41]) detected in the PNGase F-digested AD8-753EABR and TFAR-753EABR EV and VLP preparations is consistent with an Env fragment with an intact gp120-gp41 cleavage site and an adventitious cleavage site in the gp120 V3 loop. The dgp[V3+gp41] fragment comigrates with dgp120 in the gp120 Western blot shown in the upper left panel, but is evident in the gp41 Western blot shown in the lower left panel. (C) HEK 293T cells expressing the indicated Envs were incubated for 1.5 hours at room temperature with the antibodies shown. After washing, the cells were lysed and Protein A-Sepharose beads were used to precipitate antibody-Env complexes. The precipitated complexes and the cell lysates (Input) were Western blotted as described in B. (D) The supernatants of the transfected HEK 293T cells in A were incubated with TZM-bl cells. Forty-eight hours later, the TZM-bl cells were lysed and luciferase activity was measured. The means and standard deviations of technical replicates within an experiment are shown. The results are representative of those obtained in two independent experiments.

The EABR sequences dramatically increased the incorporation of the AD8 and TFAR Envs into EVs, compared to the AD8-L748STOP and TFAR-L748STOP Envs (“Env only” lanes in Fig. 3A and B). Whereas the AD8-L748STOP and TFAR-L748STOP Envs in EVs were mostly cleaved, the AD8-753EABR and TFAR-753EABR Envs in the EVs consisted of a mixture of cleaved and uncleaved Envs. Substantial fractions of the AD8-753EABR and TFAR-753EABR Envs that were uncleaved at the gp120-gp41 junction underwent another cleavage to produce fragments that migrated around 54-kDa after PNGase F digestion (labeled dgp[V3+gp41] in Fig. 3B). The dgp[V3+gp41] fragment contains both gp120 and gp41 components and likely results from adventitious proteolytic cleavage in the gp120 V3 loop (74–77). The AD8-753EABR and TFAR-753EABR Envs consisted of Endo Hf-resistant and Endo Hf-sensitive fractions, indicating that at least some of these Envs were modified by complex carbohydrates (Fig. 3B). Similar results were seen for the AD8-715EABR Env (data not shown). Even on the surface of expressing cells, the AD8-753EABR and TFAR-753EABR Envs are cleaved less than their respective AD8-L748STOP and TFAR-L748STOP Env counterparts (Fig. 3C). As expected (17–22), gp120-gp41 cleavage resulted in better recognition of the AD8-L748STOP and TFAR-L748STOP Envs by several bNAbs. By contrast, even the PTC-stabilizing changes in the TFAR-753EABR Env do not preserve the PTC, consistent with earlier suggestions that these changes stabilize the PTC much more efficiently in the context of cleaved Env (69). In summary, the large amount of uncleaved EABR-tagged Envs and the adventitious cleavage of gp120 in the EVs makes them undesirable for studies of the Env PTC.

We analyzed the particles in the supernatants of the cells expressing the L748STOP and 753EABR Envs, HIV-1 packaging proteins and an HIV-1 luciferase-expressing vector (Pseudovirus lanes in Fig. 3A and B). The AD8-753EABR and TFAR-753EABR particulate fractions included substantial amounts of Env lacking gp120-gp41 cleavage and exhibiting evidence of adventitiously cleaved gp120 (Fig. 3A and B). As these particulate fractions likely include EVs, this was expected. This fraction also includes recombinant viruses pseudotyped with the AD8-753EABR and TFAR-753EABR Envs that can infect TZM-bl cells (Fig. 3D). As expected, the EVs containing these EABR-tagged Envs did not detectably infect the TZM-bl cells in this assay; incubation of the TZM-bl cells with these EVs did not result in detectable cytotoxicity (data not shown).

In contrast to the pseudovirus preparations with the EABR-tagged Envs, the VLPs pseudotyped with the AD8-L748STOP and TFAR-L748STOP Envs contained very high percentages of cleaved Envs (Fig. 3A and B). Comparison of the pseudovirus samples digested with PNGase F and Endo Hf indicated that nearly all the AD8-L748STOP and TFAR-L748STOP Envs on the VLPs were modified by complex glycans. As previously observed (11,68,70), Env passage through the Golgi appears to be required for efficient incorporation into VLPs/virions. The pseudoviruses with the AD8-L748STOP and TFAR-L748STOP Envs infected TZM-bl cells with comparable efficiencies (Fig. 3D). Our results indicate that the proteolytic processing of the L748STOP Envs on HIV-1 VLPs is superior to that of EABR-tagged Envs on VLPs or EVs. Thus, we expect that VLPs and virions with CT truncations, when produced in permissive cells, represent a potentially useful source of mature and functional Envs, including PTC-stabilized Envs.

### Infectivity and cytopathic effects of Env CT mutants

We compared the ability of the HIV-1_AD8_ Env CT variants to mediate cell-cell fusion, virus infection and viral cytopathic effects. Cell-cell fusion mediated by the Y712STOP, L715STOP and L748STOP Envs was more efficient than that mediated by the wild-type AD8 Env or the R747L and L748A Envs (Fig. 4A). At least some of this increased syncytium-forming ability results from the increased cell-surface expression of the CT-truncated Envs.

**FIG 4.**
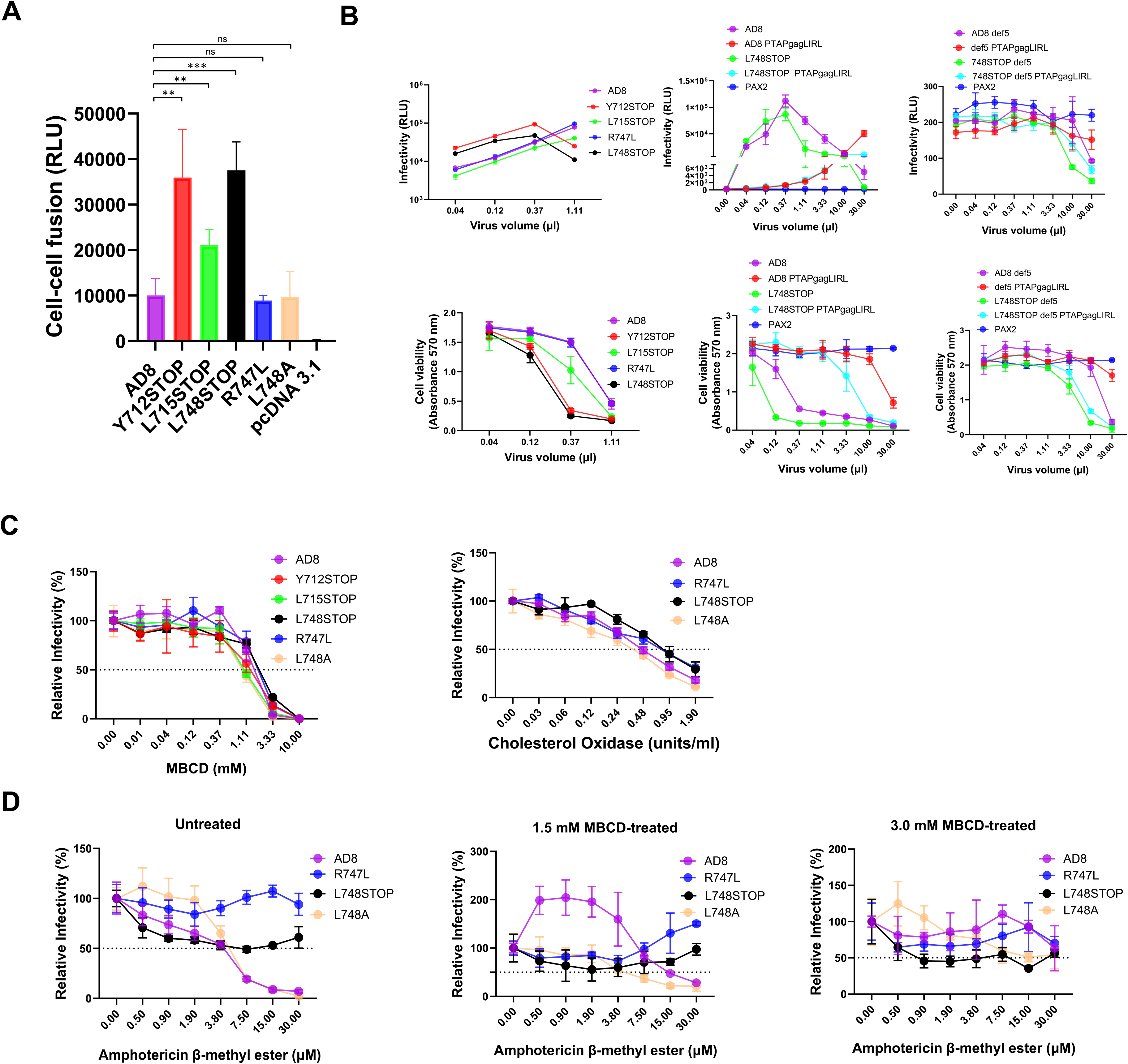
Function and inhibition of HIV-1_AD8_ Env variants. (A) The ability of the indicated HIV-1_AD8_ and mutant Env variants to mediate cell-cell fusion was measured. COS-1 effector cells transiently expressing α-gal and the indicated Env variants were cocultivated with Cf2Th-CD4/CCR5 target cells transiently expressing ω-gal at 37°C for 2 hours. Then β-galactosidase activity in the culture was measured and is reported in relative light units (RLU). The means and standard deviations derived from three independent experiments are shown. Differences between cell-cell fusion mediated by the HIV-1_AD8_ Env and that mediated by the other Envs were evaluated with an unpaired t-test (***, p<0.001; **, p<0.01; ns, not significant). (B) The infectivity and cytopathic effects of viruses with the indicated Envs were measured. Viruses were produced from HEK 293T cells transfected with pNL4-3 plasmids containing proviruses with the indicated Envs. Forty-eight hours after transfection, cell supernatants were collected and subjected to low-speed centrifugation to remove cell debris. Different volumes of the clarified cell supernatants were incubated with TZM-bl cells for 48 hours in a 37°C/5% CO_2_ incubator, after which luciferase activity in the cells was measured (upper panel). At the same time, the viability of a TZM-bl culture infected in parallel was measured. PAX2 encodes HIV-1 packaging proteins. The PTAPgagLIRL change alters a late domain in Gag (78–80,129). (C) Recombinant luciferase-expressing viruses pseudotyped with the indicated Envs were produced by transfecting HEK 293T cells with a luciferase-expressing HIV-1 vector and plasmids expressing HIV-1 packaging proteins and the indicated Envs. Forty-eight hours after transfection, viruses in the cleared cell supernatants were incubated with the indicated concentrations of methyl-β-cyclodextrin (MBCD) for 1 hour at 37°C (left panel) or cholesterol oxidase for 5 hours at 37°C (right panel). The viruses were then added to Cf2Th-CD4/CCR5 cells. After two days of culture, the cells were lysed and the luciferase activity was measured. Relative infectivity represents the luciferase activity obtained at each concentration of inhibitor compared with that observed in the absence of inhibitor. (D) Recombinant luciferase-expressing viruses pseudotyped with the indicated Envs were produced as described in C above. The pseudotyped viruses were incubated with 1.5 mM or 3 mM MBCD or control buffer for 1 hour at 37°C. The pseudotyped viruses were pelleted, resuspended and incubated with amphotericin β-methyl ester at the indicated concentrations for 1 hour at 37°C. The viruses were then added to Cf2Th-CD4/CCR5 cells. After two days of culture, the cells were lysed and luciferase activity was measured. Relative infectivity represents the luciferase activity obtained at each concentration of amphotericin compared with that observed in the absence of amphotericin. In B-D, the means and standard deviations of technical replicates within an experiment are shown. The results are representative of those obtained in at least two independent experiments.

To measure the ability of the Envs to support virus entry and cytopathic effects, IMCs encoding the Env variants were transfected into HEK 293T cells. The virus-containing cell supernatants were incubated with TZM-bl target cells. Forty-eight hours later, luciferase activity in the TZM-bl cells, as well as the cell viability, was measured. The infectivity of the Y712STOP and L748STOP viruses was higher than that of the viruses with the wild-type AD8 and other mutants Envs (Fig. 4B, upper left panel). When higher amounts of the Y712STOP and L748STOP viruses were added to the TZM-bl cells, the measured luciferase activity decreased. This decrease was related to the greater induction of cytopathic effects by the Y712STOP and L748STOP viruses (Fig. 4B, lower left panel). As expected, an alteration of the PTAP motif of the Gag polyprotein that reduces ESCRT-mediated virion release (78–80) reduced virus infectivity and cytopathic effects (Fig. 4B, middle panels). Regardless of this alteration in the Gag PTAP motif, viruses with the L748STOP Env killed the target cells more efficiently than the corresponding viruses with the wild-type AD8 Env. We wished to address whether the expression of HIV-1 proteins, including Env, in the target cells is necessary for the induction of cytopathic effects in this experimental system. To this end, we introduced the wild-type AD8 and the L748STOP *env* genes into the def5 provirus. The def5 provirus is replication-incompetent due to defective genes for reverse transcriptase, RNase H, integrase, Vif and Vpr; nonetheless, transfection of the def5 plasmid results in the production of VLPs containing Env (70). We also introduced the PTAPgagLIRL changes into the def5 proviruses. The infectivity of the def5 VLPs was very low, at the background of the assay, as expected (Fig. 4B, upper right panel). Nonetheless, VLPs with the L748STOP Envs induced greater cytopathic effects than VLPs with the wild-type AD8 Env (Fig. 4B, lower right panel). These results indicate that VLPs with the L748STOP Env can kill the TZM-bl cells even without initiating a productive infection. These results also suggest that the L748STOP Env is more cytotoxic than the wild-type AD8 Env.

### Susceptibility of viruses with Env CT variants to inhibition

Changes in the HIV-1 Env CT have been suggested to alter the sensitivity of viruses to small-molecule inhibitors and neutralizing antibodies (55,56,58–65). We examined the effect of different inhibitors on the infectivity of recombinant viruses pseudotyped by the wild-type AD8 and CT mutant Envs. We first evaluated the effects of cholesterol-modifying agents on the infectivity of the pseudotyped viruses. Methyl-β-cyclodextrin (MBCD), which depletes cholesterol from membranes (81–83), and cholesterol oxidase, which modifies membrane cholesterol (84), inhibited the viruses with the wild-type AD8 Env and CT mutant Envs comparably (Fig. 4C). Next, we investigated the susceptibility of the viruses to amphotericin B, using the less toxic amphotericin β-methyl ester (AME). We tested the hypothesis that the antiviral activity of AME depends upon cholesterol in the target membrane (60,85–90). For this purpose, we incubated the pseudotyped viruses with subneutralizing concentrations (1.5 mM or 3 mM) of MBCD or control buffer for one hour at 37°C. The viruses were then pelleted to remove MBCD, resuspended and incubated with various concentrations of AME for one hour at 37°C. The viruses were added to target cells and after two days of culture, luciferase activity in the cells was measured. For the control viruses not treated with MBCD, the viruses with the wild-type AD8 Env and L748A Env were efficiently inhibited by AME (Fig. 4D, left panel). The viruses with the R747L and L748STOP Envs were relatively resistant to AME, consistent with the proposed role of the Env CT as a determinant of amphotericin B sensitivity (59,60,90). The viruses treated with increasing concentrations of MBCD exhibited progressively greater resistance to AME (Fig. 4D, middle and right panels). The basal infectivity of the pseudotyped viruses treated with 3 mM MBCD was low, but ∼15 times the assay background; these MBCD-treated viruses were almost completely resistant to AME. These results support a model for amphotericin B antiviral activity that depends upon cholesterol in the viral membrane and upon the HIV-1 Env CT (60,90).

The sensitivity of HIV-1 infection to inhibition by specific Env ligands or by exposure to cold can provide an indication of changes in Env conformation (14,15,68,69,91–96). We examined the susceptibility of recombinant viruses pseudotyped with the HIV-1_AD8_ Env CT variants to inhibition by conformation-sensitive ligands and by cold exposure. The conformation-sensitive ligands included broadly and poorly neutralizing antibodies, sCD4-Ig, the T20 gp41 heptad repeat (HR2) region peptide, BNM-III-170 (a CD4-mimetic compound) and BMS-806. The overall patterns of sensitivity of the viruses with the wild-type AD8 and CT variant Envs to these treatments were qualitatively similar (Fig. 5A). However, in several cases, higher concentrations of antibodies/compounds were required to neutralize the viruses pseudotyped by the L712STOP, L715STOP and L748STOP Envs. Viruses with the L712STOP, L715STOP and L748STOP Envs also exhibited longer infectious half-lives after incubation on ice. Dilution of the CT-modified pseudovirus preparations reduced basal infectivity but did not change the higher bNAb IC_50_ values for the L715STOP and L748STOP pseudoviruses compared with those of the wild-type AD8 pseudoviruses (data not shown).

**FIG 5.**
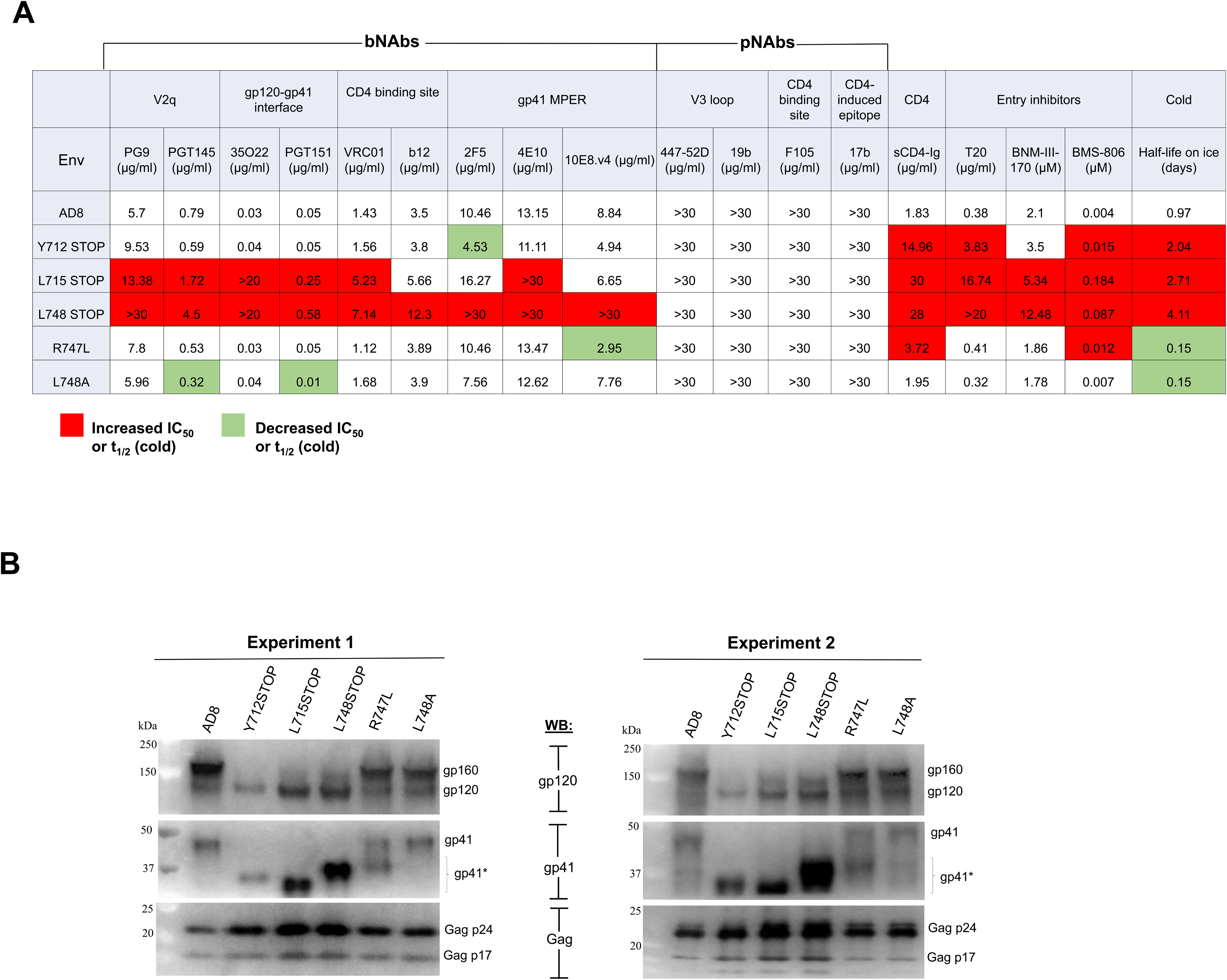
Pseudovirus Env composition and sensitivity to inhibition. (A) Pseudotyped viruses were incubated with different concentrations of antibodies/inhibitors for 1 hour at 37°C and then added to Cf2Th-CD4/CCR5 cells. After two days, the cells were lysed and luciferase activity was measured. The reported IC_50_ value of the antibody/inhibitor was calculated by fitting the data in four-parameter dose-response curves using GraphPad Prism 9. The half-life of infectivity of the pseudotyped viruses incubated on ice (0°C) was also measured. Relative to viruses with wild-type HIV-1_AD8_ Env, greater than two-fold increases (green) or decreases (red) in virus sensitivity to inhibition are indicated. The results shown are representative of those obtained in more than two independent experiments. (B) Recombinant luciferase-expressing viruses pseudotyped with the indicated Envs were produced as described in the Figure 4C legend. The pseudotyped viruses were pelleted by centrifugation at 14,000 x g for 1 hour at 4°C and then Western blotted as described in the Figure 1D legend. The results of two independent experiments are shown.

To investigate the mechanism of the relative resistance of viruses with CT-truncated Envs, we examined the composition of the pseudovirus particles. Consistent with the results shown in Fig. 1, the L715STOP and L748STOP pseudoviruses exhibited a higher level of cleaved Env than the pseudoviruses with the wild-type AD8, R747L and L748A Envs (Fig. 5B). We hypothesized that an increased level of functional Env on the viral surface results in a requirement for higher concentrations of bNAbs to neutralize the L715STOP and L748STOP pseudoviruses. To test this hypothesis, we generated pseudoviruses with lower levels of the L715STOP and L748STOP Envs by transfecting smaller amounts of the plasmids encoding these CT-truncated Envs. The level of cleaved Env (gp120/Gag p24 ratio) on the pseudoviruses was directly related to the amount of the Env-expressing plasmid transfected (Fig. 6A and Fig. 6B, left panels). The infectivity of the pseudoviruses increased and then reached a saturating level as the amount of cleaved Env on the virus increased (Fig. 6B, right panels). We tested the sensitivity of the L715STOP and L748STOP viruses with different levels of cleaved Env to neutralization by the PGT151 and 35O22 bNAbs. As the amount of cleaved Env on the L715STOP and L748STOP viruses decreased below the maximum, the IC_50_ values of both bNAbs decreased dramatically, approximating that seen for the wild-type AD8 virus (Fig. 6C). These results support the hypothesis that the number of functional, cleaved Envs on the viral surface can influence the concentration of antibody required for neutralization. The non-linear relationship between the Env content of the pseudovirus preparations and the IC_50_ values of the bNAbs has additional implications that will be addressed in the Discussion. The viruses pseudotyped by the wild-type HIV-1_AD8_ Env and the CT-modified Envs exhibited similar patterns of resistance to pNAbs and, after adjustment of virion Env levels, sensitivity to bNAbs. These observations are consistent with these functional Env variants mainly occupying the PTC.

**FIG 6.**
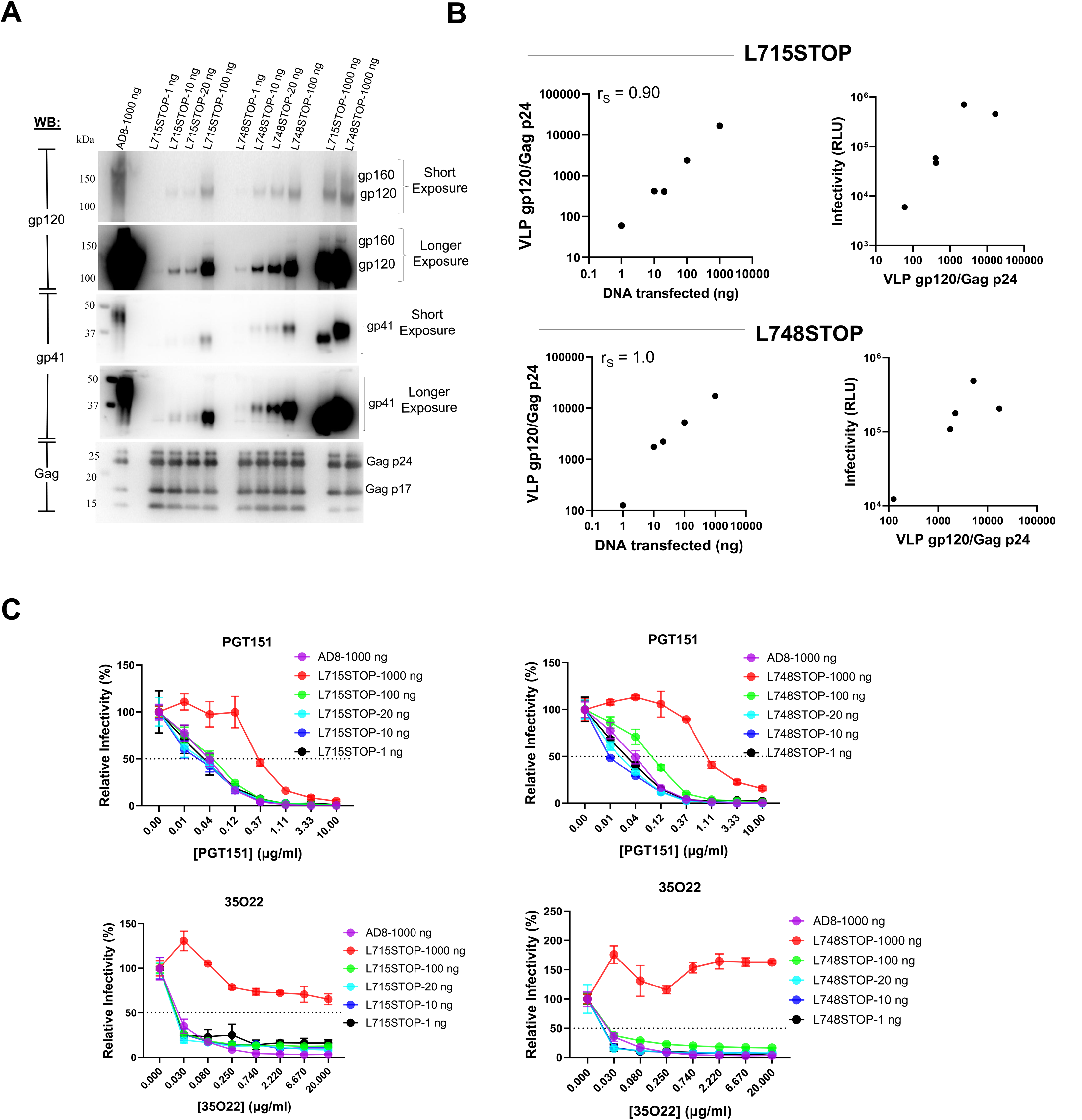
Effect of Env levels on pseudovirus infectivity and inhibition by bNAbs. (A) Recombinant luciferase-expressing viruses with different levels of the indicated Envs were produced by transfection of HEK 293T cells with a luciferase-expressing HIV-1 vector, a packaging plasmid and the indicated amounts of an Env-expressing plasmid. Forty-eight hours after transfection, the clarified cell supernatants were filtered (0.45-µm) and centrifuged at 14,000 x g for 1 hour at 4°C. The pelleted pseudoviruses were Western blotted as described in the Figure 1D legend. (B) The gp120 Env and Gag p24 bands on the Western blots were quantified using Fiji ImageJ (NIH). Infectivity was determined by adding 30 µl of the filtered supernatants from the transfected HEK 293T cells to Cf2Th-CD4/CCR5 cells and measuring the luciferase activity in the cells two days later. The panels on the left show the relationship between the amount of Env-expressing plasmid transfected and the gp120/Gag p24 ratio on the VLPs. The Spearman rank correlation coefficients (r_S_) are shown. The panels on the right show the relationship between the VLP gp120/Gag p24 ratio and the infectivity of the pseudoviruses. A parallel infection using pseudoviruses produced by transfecting 1000 ng of the wild-type HIV-1_AD8_ Env-expressing plasmid resulted in an infectivity measurement of ∼2 x 10^5^ luciferase units. (C) The neutralization of the pseudotyped viruses by the PGT151 and 35O22 bNAbs was measured as described in the legend to Figure 5A. The means and standard deviations from triplicate measurements within an experiment are shown. The experiment was repeated with comparable results.

### Effect of host restriction factors on the infectivity of HIV-1 with CT-truncated Envs

We asked if the host restriction factors SERINC5 and IFITM (97–99) inhibited the infectivity of pseudoviruses with the wild-type HIV-1_AD8_ Env and CT-truncated Envs equivalently. We produced virions by transfecting IMCs encoding the wild-type HIV-1_AD8_ Env, the L715STOP Env or the L748STOP Env with a plasmid expressing human SERINC5. In parallel negative-control experiments, we cotransfected a plasmid, ΔSERINC5, that contains a stop codon eliminating SERINC5 expression. The virions were added to TZM-bl cells, and the infectivity was evaluated by measuring luciferase activity in the cells. The viruses with the wild-type HIV-1_AD8_ Env and the CT-truncated Envs were not inhibited by SERINC5 (Fig. 7A). We also tested viruses with the laboratory-adapted HIV-1_NL4-3_ Env and its CT-truncated version NL4-3 L748STOP; these viruses were inhibited equivalently by SERINC5 (Fig. 7B), in contrast with a previous report (58). We also deleted *nef* from the IMCs producing the wild-type HIV-1_AD8_ and L748STOP Env variant and tested the viruses for sensitivity to SERINC5; the *nef* deletion did not render either virus sensitive to SERINC5 (data not shown). The expression of SERINC5 in the above studies was documented by Western blot (Fig. 7C). The wild-type HIV-1_AD8_ and L715STOP viruses were inhibited by another host restriction factor, IFITM (Fig. 7D). These results indicate that truncation of the Env CT had little effect on the inhibition of the tested HIV-1 variants by SERINC5 or IFITM.

**FIG 7.**
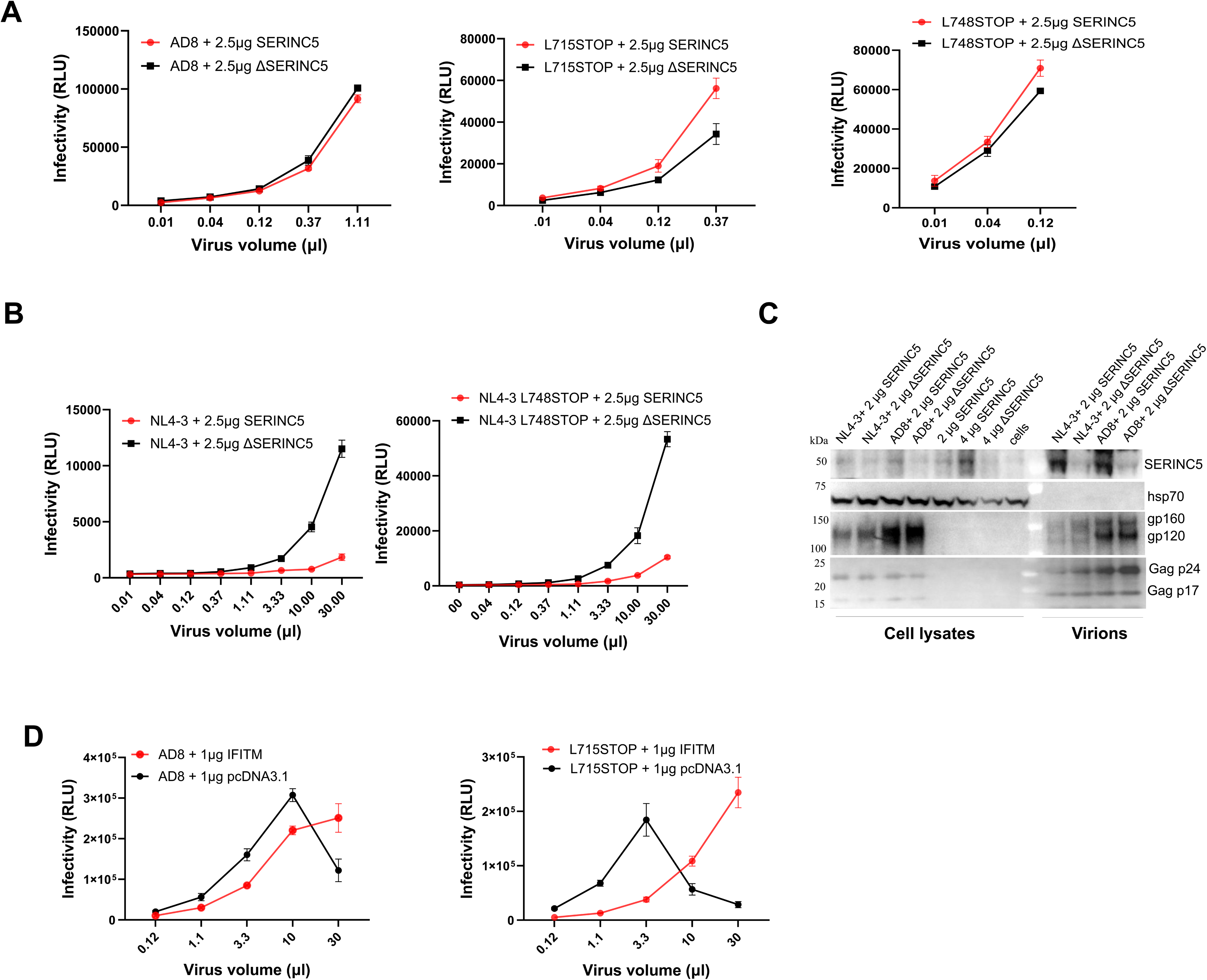
Susceptibility of pseudoviruses with Env variants to SERINC5 and IFITM restriction factors. (A) HEK 293T cells were transfected with pNL4-3 proviral constructs expressing the full-length HIV-1_AD8_ Env, the L715STOP Env or the L748STOP Env along with 2.5 µg of a plasmid encoding SERINC5 or a negative control plasmid ΔSERINC5. Forty-eight hours later, cell supernatants were cleared and incubated with TZM-bl cells. Two days later, the cells were lysed and luciferase activity was measured. (B) HEK 293T cells were cotransfected with pNL4-3 proviral plasmids encoding the wild-type HIV-1_NL4-3_ Env or the NL4-3 L748STOP Env along with 2.5 µg of the SERINC5 or ΔSERINC5 plasmids. The infectivity of the produced virions was measured on TZM-bl cells, as described in A. (C) HEK 293T cells were transfected with proviral constructs expressing the HIV-1_AD8_ and HIV-1_NL4-3_ Env variants along with the SERINC5 or ΔSERINC5 plasmids, as described in A and B above. In some experiments, the HEK 293T cells were mock transfected or transfected with the SERINC5 or ΔSERINC5 plasmids alone. Forty-eight hours after transfection, cell lysates and viruses were prepared and analyzed by Western blotting for the indicated proteins. The Western blots were developed with an anti-SERINC5 antibody (Abcam). (D) To study the effect of IFITM on HIV-1 infectivity, HEK 293T cells were transfected with pNL4-3 proviral constructs expressing the full-length HIV-1_AD8_ or L715STOP Envs along with 1 µg of a plasmid encoding IFITM or the pcDNA3.1 negative control. Forty-eight hours later, supernatants were cleared and added to TZM-bl cells. Two days later, the cells were lysed, and the luciferase activity was measured. For A-D, the results shown are typical of those obtained in at least two independent experiments.

### Inducible expression of full-length and CT-truncated Envs alone and in VLPs

To evaluate the Env phenotypes associated with CT truncation in another context, we studied A549 cell lines that inducibly express the full-length TFAR, TFAR-L715STOP and TFAR-L748STOP Envs. As mentioned above, the TFAR Env is a PTC-stabilized derivative of the HIV-1_AD8_ Env (69,70). The doxycycline-inducible polyclonal A549 cell lines were enriched for Env expression by selection with the PGT145 bNAb, which preferentially recognizes cleaved Env (66,100,101). In preliminary experiments, we found that the overall expression of the CT-truncated TFAR Envs was higher than that of the full-length TFAR Env; therefore, we used half the concentration of doxycycline to induce the cell lines expressing CT-truncated Envs as that used to induce the cell lines expressing the TFAR Env. With this adjustment, nearly equivalent levels of uncleaved and cleaved Env expression were attained in the lysates of cells expressing the TFAR, TFAR-L715STOP and TFAR-L748STOP Envs (Fig. 8A). The TFAR-L715STOP and TFAR-L748STOP Env levels on the cell surface and in EVs prepared from the cell medium were higher than those of the TFAR Env. Most of the cell surface and EV Envs were cleaved. The amount of uncleaved TFAR-L748STOP Env on the cell surface was less than that of the TFAR-L715STOP Env. Nearly all of the Envs in EVs contained complex carbohydrates (Fig. 8B). Thus, in the A549 cells, the increased levels of cell-surface and EV Envs observed for the CT-truncated Env, relative to the full-length Env, derive from Envs that have trafficked through the Golgi.

**FIG 8.**
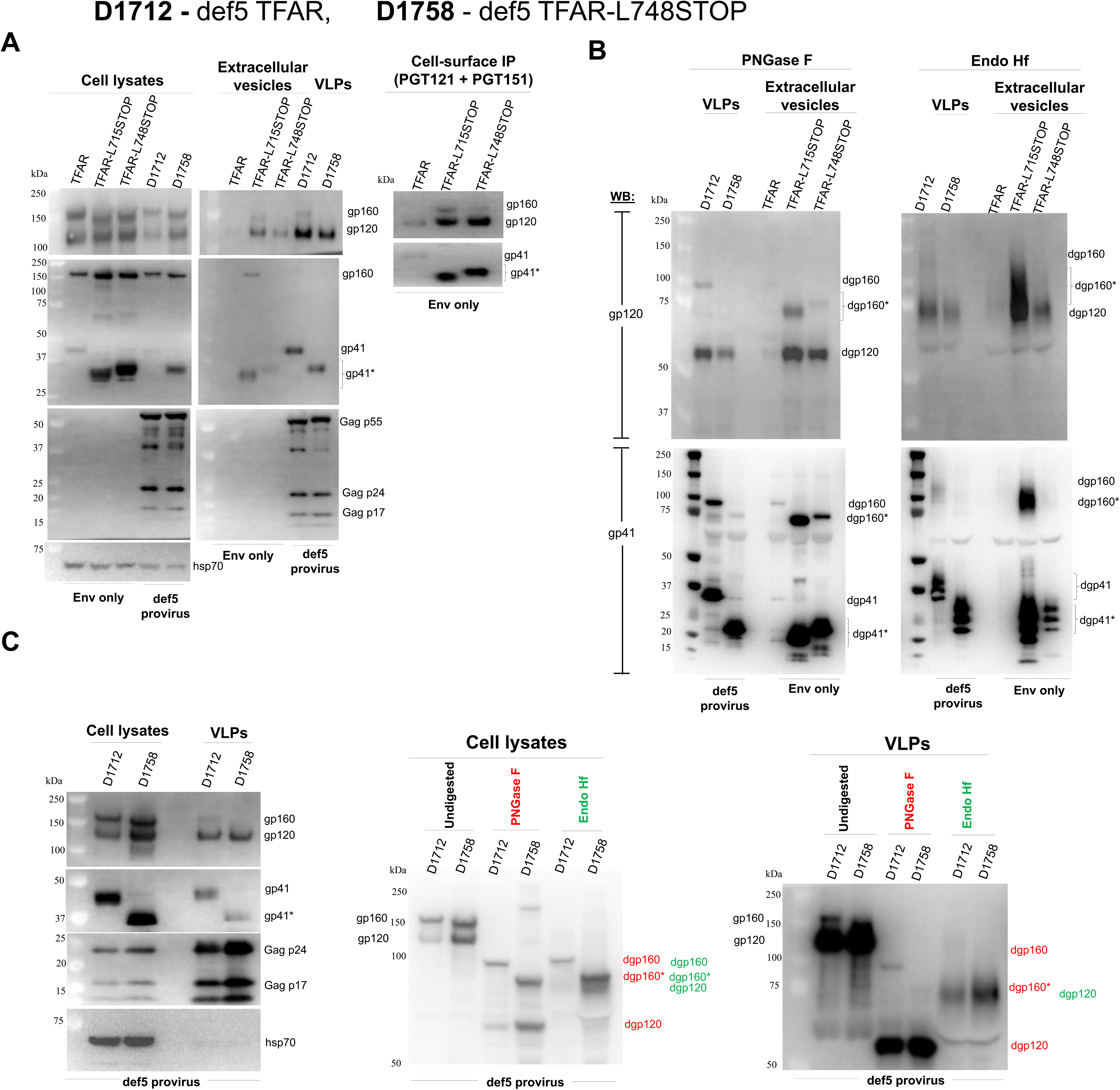
Effect of CT truncations on a PTC-stabilized Env expressed in stable, inducible cell lines. We established stable, inducible A549 cell lines expressing a PTC-stabilized Env (TFAR) and TFAR-L715STOP and TFAR-L748STOP variants. We also established A549 cells lines (designated D1712 and D1758) that inducibly express replication-defective VLPs containing the full-length TFAR Env and the TFAR-L748STOP Env, respectively. (A) These A549 cell lines were induced with 2 µg/ml doxycycline (for the full-length Envs) and 1 µg/ml doxycycline (for the CT-truncated Envs) to achieve roughly comparable Env expression in the cells. Forty-eight hours after induction, clarified cell supernatants containing VLPs were filtered (0.45-µm) and the cell supernatants containing EVs and/or VLPs were centrifuged at 14,000 x g for 1 hour at 4°C. In parallel, cell lysates were prepared. Cell-surface Envs were analyzed by incubation of intact cells with a mixture of the PGT121 and PGT151 antibodies, as described in Materials and Methods. The cell-supernatant pellets, cell lysates and immunoprecipitated cell-surface Envs were analyzed by Western blotting with a goat anti-gp120 antibody, the 4E10 anti-gp41 antibody, an anti-Gag antibody and an antibody against hsp70. (B) Extracellular vesicles and VLPs produced from the induced A549 cells in A were treated with PNGase F or Endo Hf, as described in Materials and Methods. The deglycosylated Envs were analyzed by Western blotting with a goat anti-gp120 antibody and the 4E10 anti-gp41 antibody. The results shown in panels A and B are representative of those obtained in three independent experiments. (C) To achieve comparable levels of the TFAR and TFAR-L748STOP Envs in VLPs, the D1712 and D1758 cell lines were both induced with 2 µg/ml doxycycline. Lysates of cells and VLPs were prepared and analyzed by Western blotting as in A (left panel). The samples were also digested with PNGase F and Endo Hf and analyzed by Western blotting with a goat anti-gp120 antibody (middle and right panels). The Env bands resulting from PNGase F treatment are labeled red and the Env bands from Endo Hf treatment are labeled green.

We also studied A549 cell lines that inducibly express replication-defective def5 HIV-1 proviruses with the TFAR and TFAR-L748STOP Envs; these cell lines, which produce defective VLPs, are designated D1712 (with the TFAR Env) and D1758 (with the TFAR-L748STOP Env) (70). Both cell lines were induced with 2 µg/ml doxycycline. The total levels of TFAR Env and TFAR-L748STOP Env expressed in the cell lysates and VLPs were similar (Fig. 8A). The two Envs were also processed comparably. Nearly all the TFAR and TFAR-L748STOP Envs in the VLPs contained complex glycans (Fig. 8B and C). The L748STOP truncation resulted in decreased incorporation of the uncleaved Env into the VLPs (Fig. 8A and C). Thus, even in VLPs produced from HIV-1 proviruses, which generally exhibit high ratios of cleaved:uncleaved Env (11,68,70), the L748STOP truncation can help to decrease the amount of uncleaved Env.

We compared the antigenic profile of the TFAR and TFAR-L748STOP Envs on the surface of the A549 cells transduced with def5 proviruses and on VLPs (Fig. 9). On the cell surface, both cleaved Envs were recognized by bNAbs but not by pNAbs, as expected for Envs in a PTC. The cleaved TFAR-L748STOP Env on the cell surface bound the b12 CD4-binding site (CD4BS) bNAb better than the cleaved TFAR Env; another CD4BS bNAb, 3BNC117, bound the TFAR-L748STOP Env slightly worse than the TFAR Env (Fig. 9A and B, left panels). The b12 bNAb has been suggested to bind an occluded intermediate conformation distinct from the PTC, which is recognized by other CD4BS bNAbs (102–107). The antigenic profiles of the TFAR and TFAR-L748STOP Envs on VLPs were indistinguishable (Fig. 9A and B, right panels). Thus, the VLPs produced by the D1758 cell line represent a source of mature, PTC-stabilized HIV-1 Env for further study.

**FIG 9.**
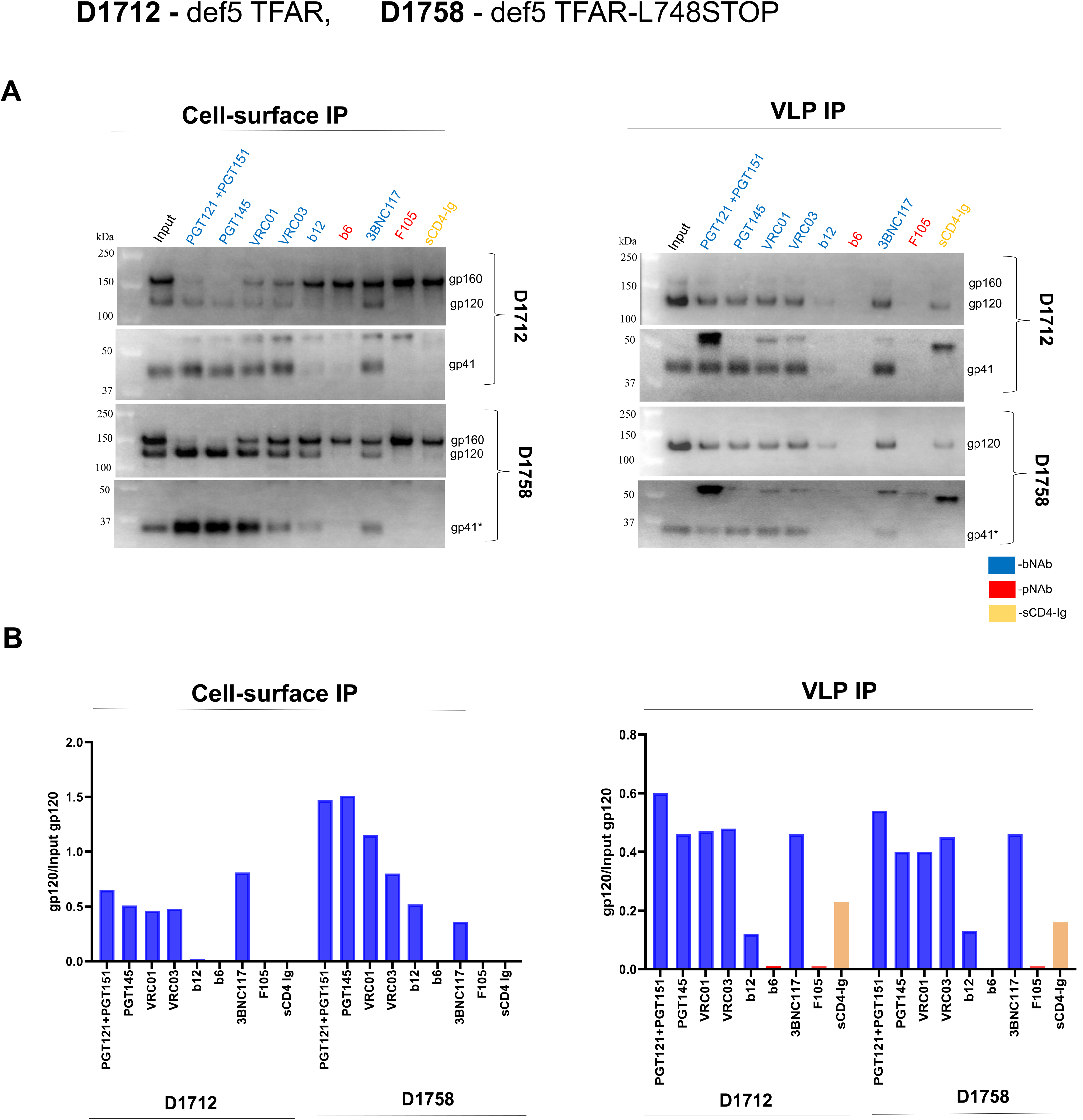
Antigenic profile of the cell-surface and VLP Envs. The d1712 and D1758 cell lines expressing defective VLPs with the full-length TFAR Env and TFAR-L748STOP Env were induced with 2 µg/ml doxycycline. Forty-eight hours later, VLPs were prepared from the cell supernatants as described in the Figure 6 legend. To serve as input samples, a portion of the cells and VLPs was lysed in detergent. The main portion of the cells (for the cell-surface immunoprecipitation (IP)) and VLPs (for the VLP IP) was incubated with a panel of antibodies or sCD4-Ig for 1.5 hours at room temperature. The cells were washed with 1x PBS. The VLPs were pelleted and washed with 1x PBS. The cells and VLPs were then lysed and the lysates incubated with Protein A-agarose beads for 1 hour at 37°C. The beads were washed three times. The precipitated proteins and input samples were Western blotted with a goat anti-gp120 antibody and the 4E10 anti-gp41 antibody. Quantification of the gp120 bands was performed in Fiji ImageJ (NIH). The ratio of precipitated gp120 to the input gp120 is reported in the bar graphs. The names of the Env ligands and the bars on the graphs are colored as follows: broadly neutralizing antibodies (bNAbs) – blue; poorly neutralizing antibodies (pNAbs) – red; sCD4-Ig – yellow ochre. The results of a typical experiment are shown. The experiment was repeated with similar results.

### Phenotypes of CT truncation in Envs of other HIV-1 strains

To determine if the phenotypes observed for the HIV-1_AD8_ L748STOP Env hold true for other primary HIV-1 Envs, we introduced this change into the Envs of six HIV-1 strains from multiple clades. To evaluate Env expression and processing, the Env expressor plasmids were transfected into HEK 293T cells along with the psPAX2 plasmid expressing HIV-1 packaging proteins. The cell lysates and VLPs were Western blotted (Fig. 10A). The expression levels of the full-length Envs and their L748STOP Env counterparts in cell lysates were comparable. As seen in the case of the HIV-1_AD8_ Env, different gp41 glycoforms were observed. Unlike the case for the HIV-1_AD8_ Env, the L748STOP truncation of the CT did not increase the levels on VLPs of Envs from other HIV-1 strains. However, in every HIV-1 strain examined, the cleaved:uncleaved Env ratio on VLPs was greater for the L748STOP Env compared with the full-length Env. Thus, in multiple diverse HIV-1 strains, the L748STOP CT truncation can enrich the proportion of mature Env in pseudovirus VLP preparations.

**FIG 10.**
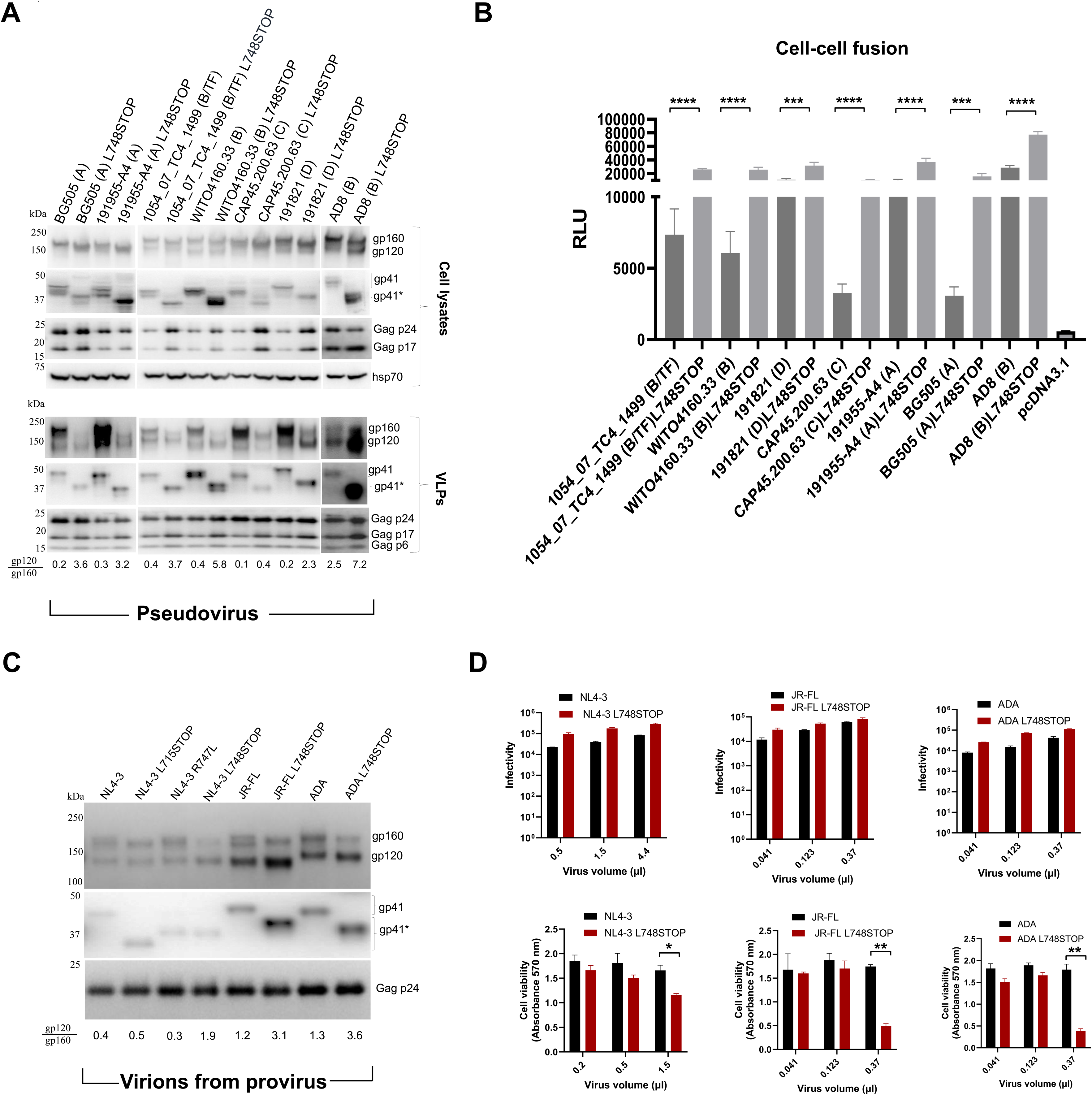
Effect of CT truncation on Envs of diverse HIV-1 strains. (A) HEK 293T cells were transfected with plasmids expressing the full-length and L748STOP Envs derived from the indicated HIV-1 strains; the phylogenetic clade and transmitted/founder (TF) status of the viruses are indicated in parentheses. The Env-expressing plasmids were cotransfected with an HIV-1 luciferase-expressing vector and the psPAX2 plasmid that expresses HIV-1 packaging proteins. Forty-eight hours later, the cell supernatants were collected, filtered through a 0.45-µm membrane, and centrifuged at 14,000 x g for 1 hour at 4°C. In parallel, the cells were lysed. The pelleted pseudovirus VLPs and clarified cell lysates were Western blotted to detect the indicated proteins. The gp120/gp160 ratio in the VLPs is shown beneath the Western blot. (B) The ability of the full-length and L748STOP Envs from different HIV-1 strains to mediate cell-cell fusion was tested. COS-1 effector cells transiently expressing α-gal and the indicated Envs were cocultivated with Cf2Th-CD4/CCR5 target cells transiently expressing ω-gal for 2 hours at 37°C. Then the β-galactosidase activity in the cells was measured. The results shown are the means and standard deviations derived from three independent experiments. The differences in cell-cell fusion between the full-length and L748STOP Envs were evaluated with an unpaired t-test (****, p<0.0001; ***, p<0.001; ns, not significant). (C) HEK 293T cells were transfected with infectious provirus constructs encoding the indicated Envs. Forty-eight hours later, virions were concentrated from cell supernatants and processed as described in A. (D) HEK 293T cells were transfected with proviral plasmids encoding the full-length and L748STOP Envs from the indicated HIV-1 strains. Forty-eight hours later, virus-containing cell supernatants were centrifuged at low speed to remove cell debris. Different volumes of the virus preparations were added to TZM-bl target cells. One set of the TZM-bl target cells was harvested two days later to measure luciferase activity, an indicator of virus infectivity (upper panels). The viability of a second set of TZM-bl target cells was measured 72 hours after infection (lower panel). The means and standard errors of the mean of triplicate measurements derived from a representative experiment are reported. The statistical significance of the observed differences was evaluated with an unpaired t-test. Significant differences are indicated (**, p<0.01; *, p<0.05).

As was seen for the HIV-1_AD8_ Env, the Envs of primary HIV-1 strains mediate cell-cell fusion more efficiently when the CT is truncated by the L748STOP change (Fig. 10B).

We studied the phenotypes resulting from CT modifications of Envs on virions produced by IMCs (Fig. 10C). The L748STOP change was introduced into the Envs encoded by the HIV-1_NL4-3_, HIV-1_JR_-_FL_ and HIV-1_ADA_ Env IMCs. The HIV-1_ADA_ Env is derived from the same virus isolate as the HIV-1_AD8_ Env and these Envs are thus related, but not identical (108,109). In addition, the L715STOP and R747L changes were introduced into the HIV-1_NL4-3_ Env encoded by the IMC. The IMCs were transfected into HEK 293T cells and the virions produced in the cell medium were analyzed. The full-length HIV-1_NL4-3_ Env and CT variants were expressed at similar levels on virions. The HIV-1_NL4-3_ R747L Env was clipped in the gp41 CT, as expected (60). Compared with the other HIV-1_NL4-3_ Env variants, the HIV-1_NL4-3_ L748STOP Env was processed efficiently and gp160 was relatively excluded from virions. The cleaved:uncleaved ratios of the HIV-1_JR_-_FL_ L748STOP and HIV-1_ADA_ L748STOP Envs on virions were increased compared with those of the respective full-length Envs. These results confirm that the L748STOP CT truncation can reduce the relative level of uncleaved Env on virion preparations from multiple HIV-1 strains.

We analyzed the infectivity and cytotoxicity of the virions with HIV-1_NL4-3_, HIV-1_JR_-_FL_ and HIV-1_ADA_ full-length Envs and the L748STOP Envs (Fig. 10D). The L748STOP Envs supported infection of TZM-bl cells as well or better than the respective full-length Envs. At the highest dose of virus tested, the virions with the 748STOP Env were more cytotoxic to the TZM-bl cells than virions with the respective full-length Envs. Thus, as was seen for the provirus-produced virions with the HIV-1_AD8_ Env, CT truncation resulting from the L748STOP change results in highly infectious, cytotoxic virions.

## DISCUSSION

The identification of homogeneous sources of the PTC of membrane HIV-1 Envs would assist studies of the structure and immunogenicity of this metastable state. In cells producing viruses or VLPs, HIV-1 Envs are transported to the cell surface by two pathways (11) (Fig. 11). In the conventional secretory pathway, Envs in the Golgi are modified by complex glycans and cleaved on their way to the cell surface. Golgi-processed Envs are preferentially incorporated into virions via a process that, in non-permissive cells, is dependent upon the CT (23,24,39–49). A fraction of Env bypasses the Golgi and is transported to the cell surface in a form that is uncleaved and lacks complex carbohydrates (11). Although largely excluded from virions, these Envs can be incorporated into EVs, a common contaminant of virion/VLP preparations. Envs that are uncleaved as a result of either bypassing the Golgi or overwhelming the Golgi furin protease are conformationally flexible and recognized by pNAbs (17–22,110). Therefore, minimizing or eliminating uncleaved Env in preparations of virions/VLPs would enhance Env PTC homogeneity.

**FIG 11.**
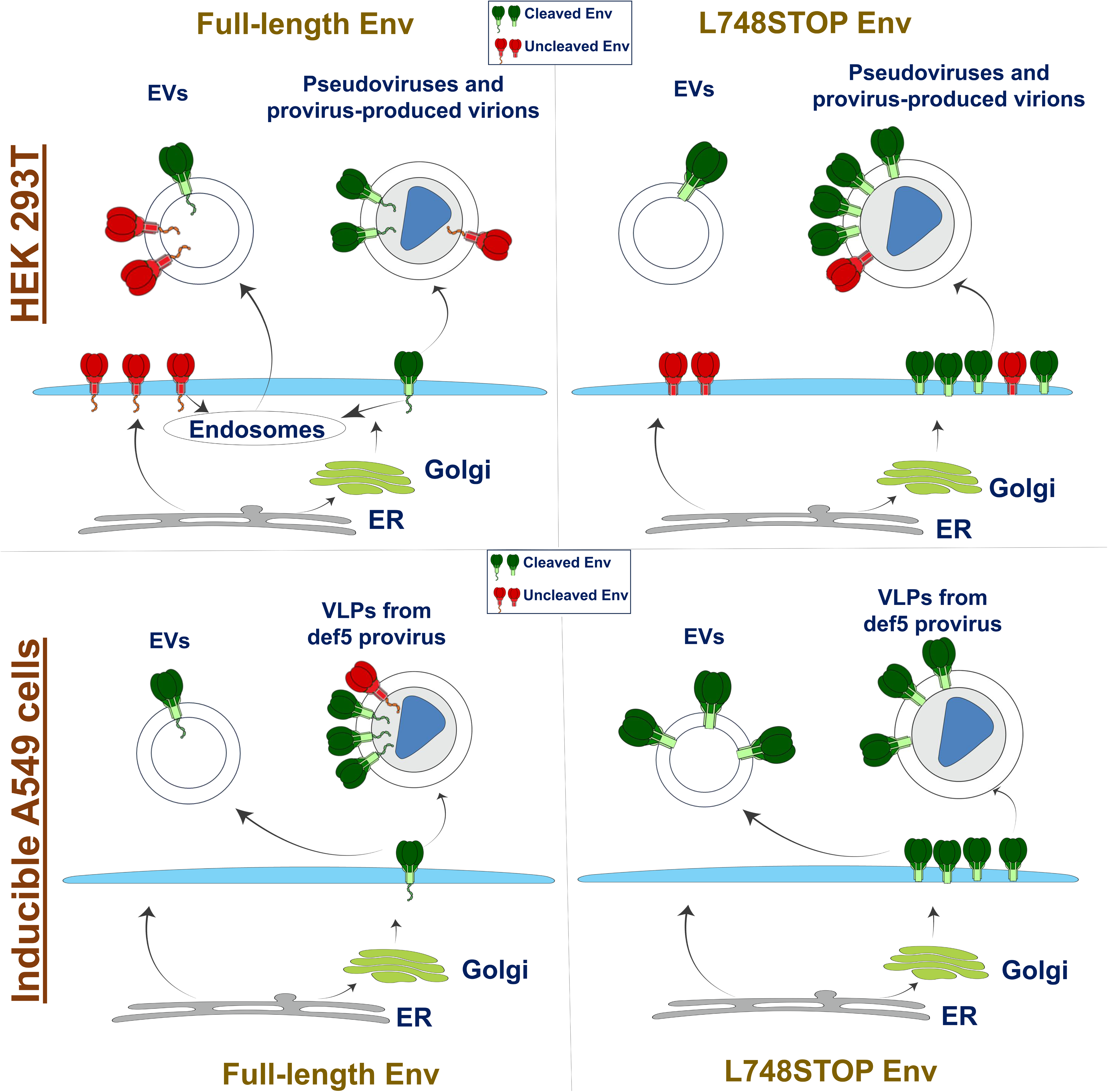
Summary of the effects of CT truncation on Env expression on cell surfaces, virions and extracellular vesicles. Two pathways by which HIV-1 Envs synthesized in the endoplasmic reticulum (ER) traffic to the cell membrane are depicted: 1) Envs in the conventional secretory pathway pass through the Golgi complex, where most Envs are cleaved, modified by complex glycans, and potentially able to be incorporated into virions; and 2) Envs in the unconventional secretory pathway bypass the Golgi, are uncleaved, and are largely excluded from virions (11). The effects of CT truncation (L748STOP) on the levels and cleavage of Envs on cell surfaces, extracellular vesicles (EVs) and pseudovirus VLPs and virions are illustrated. Cleaved Envs are shown in green and uncleaved Envs in red. The CT is depicted on the full-length Envs. The results obtained in transiently expressing HEK 293T cells and in stably expressing A549 cells are summarized.

We investigated the potential effect of modification of the Env CT on the incorporation of cleaved and uncleaved Envs into virions, VLPs and EVs in permissive cells. The CT was completely removed from the Y712STOP and L715STOP Envs. For some natural HIV-1 Env variants, partial clipping of the Env CT by the viral protease occurs (60). Based on the site in the CT at which this clipping occurs, we studied the effects of a corresponding Env truncation (L748STOP) on Env phenotypes (Fig. 11). Compared with the full-length Env, in permissive HEK 293T cells, the CT truncations resulted in higher Env levels on the cell surface and increased ratios of cleaved:uncleaved Env in VLPs pseudotyped with Envs from multiple HIV-1 strains. In virions produced in HEK 293T cells from proviral IMCs, in which Env processing is generally efficient (68,70), CT truncation allowed efficient virion Env incorporation while maintaining a high cleaved:uncleaved Env ratio. Although the different CT-truncated Envs shared many phenotypes, we found that the L748STOP mutant consistently exhibited efficient expression and processing and therefore included this mutant in most of the experiments. In HEK 293T cells, the high levels of cleaved L748STOP Env on the cell surface are apparently available for incorporation into virions and VLPs, but are not efficiently incorporated into EVs (Fig. 11). In these cells, the endocytosis signals on the CT likely influence the efficiency with which cell-surface Envs are recycled into endosomes/multivesicular bodies (23,24,33–37). Exosomes, a major class of EVs, are generated by inward budding of the late endosomes/multivesicular bodies (111–114). At diameters of 30-150 nm, exosomes overlap HIV-1 virions in size and thus are frequent contaminants of virion/VLP preparations (70). Whereas the full-length Env in EVs exhibits more uncleaved than cleaved Envs, resembling the proportion on the cell surface, the small amount of the L748STOP Env in EVs is completely cleaved. Therefore, contaminating EVs contribute less uncleaved Env to VLP preparations in the case of the L748STOP Env. For both full-length and CT-truncated Envs, the cleaved Envs on virions/VLPs are modified by complex glycans. In preparations of VLPs produced in HEK 293T cells, a fraction of the full-length uncleaved Env is Endo Hf-sensitive, suggesting that these Envs may have bypassed the Golgi. On the other hand, the uncleaved L748STOP Env in VLPs is Endo Hf-resistant, consistent with passage through the Golgi. Thus, compared with the full-length Env, a higher proportion of the L748STOP Env appears to traffic through the Golgi, populate the cell surface and become incorporated into virions/VLPs. At the same time, cell-surface L748STOP Envs, some of which are uncleaved, avoid endocytosis and eventual incorporation into exosomes/EVs (Fig. 11).

We recently reported the inducible production from A549 cells of replication-defective VLPs containing Envs stabilized in a PTC (70). To evaluate whether the observations made in HEK 293T cells might apply to another, more practical VLP production system, we compared A549 cells producing VLPs with either the full-length or L748STOP PTC-stabilized TFAR Env. During their establishment (70), the A549 cell lines had been enriched for cell-surface expression of cleaved Env by selection with the PGT145 bNAb, which preferentially recognizes the mature Env (100,101). As in the HEK 293T cells, compared to the full-length TFAR Env, the TFAR L748STOP Env exhibited a higher overall level of expression and much higher cell-surface expression of cleaved Envs (Fig. 11). Perhaps because of the selection with the PGT145 bNAb, nearly all the full-length and L748STOP TFAR Envs on VLPs and EVs were cleaved. A small amount of Endo Hf-resistant full-length TFAR Env on VLPs was uncleaved; relatively little uncleaved TFAR L748STOP Env was detected on VLPs but was also Endo Hf-resistant. These low levels of uncleaved Env have apparently passed through the Golgi and acquired some complex glycans but escaped furin processing. More cleaved TFAR L748STOP Env was observed on EVs compared with the full-length TFAR Env. Unlike the case in HEK 293T cells, in A549 cells, the CT does not appear to be important for efficient Env incorporation into EVs. The levels of full-length Env and L748STOP Env in the EVs from A549 cells reflect the levels of those Envs on the cell surface; these EVs may represent microvesicles, which bud directly from the plasma membrane (111–114). In summary, even in cells selected for expression of cleaved HIV-1 Envs, the L748STOP truncation can decrease the small amount of uncleaved Env incorporated into VLP preparations.

We also evaluated HIV-1 Env-EABR fusion proteins as a means of producing EVs that efficiently incorporate oriented Env trimers, an approach that produced immunogenic SARS-CoV-2 S glycoprotein EVs (73). Unfortunately, a substantial fraction of the HIV-1 Env-EABR fusion proteins was not proteolytically cleaved at the gp120-gp41 junction. This was the case regardless of the placement of the EABR sequence at residue 715 or 753 of the CT, or whether the wild-type HIV-1_AD8_ Env or the more efficiently processed TFAR Env was used. Adventitious cleavage of the HIV-1 Env-EABR fusion proteins in gp120, presumably in the V3 loop (74–77), further suggested that the Env components of these EVs are conformationally heterogeneous. The cell-surface Env-EABR fusion proteins were largely uncleaved and recognized by pNAbs but not the PGT145 bNAb. At least a fraction of the HIV-1 Env-EABR fusion proteins is functional, as these proteins supported the entry of a luciferase-expressing pseudovirus. Based on these results, HIV-1 VLP preparations containing the CT-truncated Envs appear to be a much better source of homogeneous, cleaved, and PTC-stabilized Env than the Env-EABR EVs.

Preserving the native structure of membrane HIV-1 Env in VLP preparations is important to the relevance and reliability of studies utilizing these particles. As some truncations of the CT have been reported to affect the conformation of the Env ectodomain (55,56), we evaluated the susceptibility to inhibition of recombinant viruses with full-length and CT-truncated Envs. Env CT truncations had no significant effect on the susceptibility of the tested HIV-1 strains to the host restriction factors SERINC5 and IFITM, a result apparently at odds with those reported in reference 58. Viruses with the L715STOP and L748STOP Envs exhibited increases in the IC_50_ values for multiple bNAbs, sCD4-Ig, T20, the CD4mc BNM-III-170, and BMS-806. However, viruses with these CT-truncated Envs remained resistant to neutralization by pNAbs. Thus, although the qualitative pattern of sensitivity is similar for the viruses with the wild-type HIV-1_AD8_ and CT-truncated Envs, quantitative differences in the IC_50_ values of multiple ligands resulted from the increased concentrations of CT-truncated Envs on virions. When the virion levels of the L715STOP and L748STOP Envs were reduced, the bNAb IC_50_ values were similar for viruses with the wild-type HIV-1_AD8_ Env and the CT-truncated Envs. These results suggest that the wild-type HIV-1_AD8_ Env and the Envs with CT truncations beginning at residues 715 and 748 have similar ectodomain conformations.

The observed relationship between virion Env levels and bNAb concentrations required for neutralization provides information relevant to studies of HIV-1 entry stoichiometry. Definition of T, the number of Env trimers required for entry, is complicated by the low infectivity:particle ratio of HIV-1 preparations (115–118). This leads to uncertainties about the functional virion Env number, distribution, mobility and ability to participate in productive entry events; all of these parameters can potentially influence the estimation of T (119–124). We encompass these multiple parameters in the variable n_t_, which describes the number of functional Env trimers on a virus particle that can participate in the virus entry process. On an infectious virion, n_t_ must be greater than or equal to T. The highest level of CT-truncated Envs on virions achieved in our experimental system was beyond the Env level required to saturate virus infectivity; in these cases, n_t_ must be greater than or equal to T+1. This redundancy in the amount of virion Env needed for entry is reflected in the increased amount of bNAb required to inhibit virus entry. At lower Env levels on the virions, over a 40-fold range of gp120:Gag p24 ratios, there is a linear relationship between the virion Env levels and infectivity. In this linear range, infectivity is limited by n_t,_ implying that n_t_ is very close to T for most virions. Over this wide range of virion Env levels, the neutralization of viruses with the wild-type HIV-1_AD8_ Env and CT-truncated Envs is comparable. In this experimental system, the results are best explained by a model in which: 1) Env is distributed among the virus particles randomly; 2) For most of the range of Env levels per virion achieved, n_t_ is less than or equal to T; and 3) Only at the highest levels of Env achieved is there redundancy in the entry process, where n_t_ is greater than T+1. Thus, even in systems using CT-truncated Envs, n_t_ is only marginally greater than T; this result should inform estimates of n_t_ made in systems using full-length Envs, where the virion Env concentrations are typically lower. Indeed, several studies suggesting low values of T have been based on estimates of n_t_ that are equal to or only slightly greater than T (118,121,122).

We evaluated the infectivity and induction of cytopathic effects by recombinant viruses with full-length and CT-truncated Envs. The relative increases in the levels of CT-truncated Envs on the surface of expressing cells was associated with increased syncytium-forming ability. The relative increases in the levels of CT-truncated Envs on virions were associated with increased infectivity and induction of cytopathic effects in the target cells. This cytotoxicity was retained by the replication-defective def5 viruses, and therefore did not require infection of the target cells. Changes in the Gag PTAP motif significantly reduced the cytotoxicity, suggesting that VLPs mediate the cytopathic effects. Incubation of CD4+/CCR5+ cells with sufficiently high concentrations of virions results in “fusion from without”, a cytotoxic process that has been reported to be enhanced by deletion of the Env CT (125,126).

In contrast to the CT truncations, the CT changes (R747L and L748A) near the viral protease cleavage site exhibited limited phenotypes. As this proteolytic clipping of the CT occurs only after the HIV-1 protease dimerizes and becomes active during virus assembly and release, these changes do not influence Env intracellular transport, processing, cell-surface expression or virion incorporation. With one exception, increases (R747L) and decreases (L748A) in the degree of protease clipping of gp41 in pseudovirus VLPs exhibited no detectable phenotype. This implies that the major phenotypes associated with the L748STOP Env arise because of effects on one or more of the above processes, rather than an effect of the truncated virion Env CT *per se*. The one phenotype associated with the truncated Envs in the R747L and L748STOP mutant virions is resistance to amphotericin β-methyl ester (AME). Our results suggest that both the Env CT and cholesterol contribute to the antiviral activity of AME, supporting models in which AME and cholesterol create pores in the viral membrane (59,60,90). A cholesterol-binding element has been mapped to the HIV-1 gp41 CT (127).

These studies can guide efforts to utilize more homogeneous sources of HIV-1 Envs to characterize the structure and immunogenicity of the elusive PTC.

## MATERIALS AND METHODS

### Plasmids

The full-length HIV-1_AD8_ Env and CT-modified Envs were expressed using the pSVIIIenv plasmid. The *env* genes encoding the L715STOP and L748STOP Envs in the pSVIIIenv plasmids have stop codons replacing the codons for Leu 715 and Leu 748, respectively. The Y712STOP Env encoded by the pSVIIIenv plasmid has a carboxy-terminal Gly_3_His_6_ tag; in the *env* gene encoding the Y712 Env, a stop codon replaces the codon for Tyr 712. The primary HIV-1 Envs [(191955-A4 (Clade A), BG505 (Clade A), 1054_07_TC4_1499 (Clade B/TF), WITO4160.33 (Clade B), CAP45.200.63 (Clade C) and 191821 (Clade D)] were expressed using pcDNA3.1 (117). For experiments in which the Envs were expressed in the context of an HIV-1 provirus, the *env* genes were cloned into the pNL4-3 IMC, obtained from the NIH HIV Reagent Program (128). The *env* genes encoding the Y712STOP, L715STOP and L748STOP Envs in the pNL4-3 IMCs have stop codons replacing the codons for Tyr 712, Leu 715 and Leu 748, respectively. The construction of the JR-FL IMC was described in reference 68. The AD8 and ADA *env* genes were cloned into the pNL4-3 proviral plasmid between 5’ cagaagacagtggca and 3’gatgggtggcaagtg sequences. Def5 was created by introducing multiple stop codons into the AD8 IMC that eliminate the expression of reverse transcriptase (RT), RNase H, integrase (IN), Vif, and Vpr (70). The codon-optimized sequence encoding EABR (FNSSINNIHEMEIQLKDALEKNQQWLVYDQQREVYVKGLLAKIFELEKKTETAAHSLP) followed by two stop codons was inserted after the codon for Ser 752 in the pSVIIIenv plasmids expressing the HIV-1_AD8_ and TFAR Envs. The EABR sequence was inserted after the codon for Pro 714 to make the pSVIIIenv plasmid expressing the AD8-715EABR Env.

The PTAP motif within the p6 domain of the HIV-1 Gag protein (78–80,129) was changed to LIRL using Q5 site-directed mutagenesis (New England BioLabs). The pBJ5 plasmid expressing SERINC5 was generously provided by Dr. Heinrich Göttlinger (UMass Worcester) (98). In the control ΔSERINC5 plasmid, two stop codons were introduced by mutating the initiation codon ATG of SERINC5. The pQCXIP plasmid expressing IFITM1 was obtained from Addgene.

The DNA sequences of all constructs were confirmed.

### Cell lines

HEK 293T, TZM-bl and A549 cell lines were cultured in Dulbecco’s modified Eagles’ medium (DMEM) supplemented with 10% fetal bovine serum (FBS) and 100 µg/mL penicillin-streptomycin (Life Technologies). Cf2Th-CD4/CCR5 cells stably expressing the human CD4 and CCR5 coreceptors for HIV-1 were grown in the same medium supplemented with 0.4 mg/ml of G418 and 0.2 mg/ml of hygromycin.

A549 cell lines inducibly expressing the HIV-1 Rev protein and the full-length TFAR, TFAR-L715STOP and TFAR-L748STOP Envs were established as previously described (130). The TFAR Env is a PTC-stabilized derivative of the HIV-1_AD8_ Env (69).

The D1712 and D1758 A549 cell lines contain the doxycycline-inducible def5 provirus with the TFAR and TFAR-L748STOP Envs, respectively, and were established as described (70). The def5 proviruses are defective in the expression of reverse transcriptase, RNAse H, integrase, Vif and Vpr. In addition, the transactivation response (TAR) element in the def5 long terminal repeat (LTR) is mutated to reduce Tat-mediated transactivation. Two tet operator sequences have been inserted into the U3 regions of the LTRs to upregulate transcription in the presence of doxycycline. The D1712 and D1758 cells were maintained as polyclonal cell lines.

### Small-molecule compounds

The CD4-mimetic compound (CD4mc) BNM-III-170 was synthesized as described previously (37,57). The compound was dissolved in dimethyl sulfoxide (DMSO) at a stock concentration of 10 mM and diluted to the appropriate concentration in cell culture medium for antiviral assays. BMS-806 was purchased from Selleckchem. Ritonavir and methyl-β-cyclodextrin was purchased from Sigma-Aldrich and amphotericin β-methyl ester from Santa Cruz Biotechnology.

### Enzymes

Protein N-glycosidase F (PNGase F) and endoglycosidase Hf (Endo Hf) were purchased from New England BioLabs. Cholesterol oxidase was purchased from Sigma-Aldrich.

### Antibodies

Poorly neutralizing antibodies (F105, 19b, 447-52D, 17b and F240) and broadly neutralizing antibodies (VRC01, VRC03, b12, b6, PGT145, PG9, PGT151, 35O22, PGT121, 3BNC117, 2F5, 4E10 and 10E8.v4) against the HIV-1 Env were obtained through the NIH HIV Reagent Program, Division of AIDS, NIAID, NIH. The bNAbs included PGT121 against V3 glycans; VRC03, VRC01 and b12 against the gp120 CD4-binding site (CD4BS); PG9 and PGT145 against quaternary V2 epitopes at the Env trimer apex; PGT151 and 35O22 against the gp120-gp41 interface; and 2F5, 4E10 and 10E8.v4 against the gp41 membrane-proximal external region (MPER). The pNAbs included F105 against the gp120 CD4BS; 19b, 39F and 447-52D against the gp120 V3 region; 17b against gp120 CD4i epitopes; and F240 against a Cluster I epitope on gp41. Western blots were developed with 1:2,000 goat anti-gp120 polyclonal antibody (Invitrogen), 1:2,000 4E10 anti-gp41 antibody, 1:3,000 rabbit anti-gag (p55/p24/p17) antibody (Abcam), 1:10,000 rabbit anti-hsp70 (K-20) antibody (Santa Cruz Biotechnology), 1:250 rabbit anti-SERINC5 antibody (Abcam). The HRP-conjugated secondary antibodies were 1:2,000 rabbit anti-goat (Invitrogen), 1:2,000 goat anti-human (Invitrogen), and 1:10,000 anti-rabbit (Sigma-Aldrich).

### Expression and processing of HIV-1 Env variants

HEK 293T cells were transfected transiently either with: 1) pSVIIIenv plasmids encoding HIV-1 Env and Rev along with a plasmid encoding HIV-1 Tat at an 8:1 ratio; or 2) pcDNA3.1 plasmids encoding Env. In some experiments, the psPAX2 plasmid encoding HIV-1 packaging proteins and a luciferase-expressing HIV-1 vector were transfected with the Env-expressing plasmids (94,96). Forty-eight hours later, cell lysates were prepared in 1x PBS/0.5% NP-40/protease inhibitors. Extracellular vesicles (EVs) were prepared from the medium of Env-expressing cells; the cell supernatant was cleared (600 x g for 10 min) followed by centrifugation at 14,000 x g for 1 hour at 4°C. In experiments in which pseudotyped VLPs or provirus-produced virions were studied, HEK 293T cell supernatants were cleared (600 x g for 10 min), filtered through a 0.45-µm membrane, and centrifuged at 14,000 x g for 1 hour at 4°C.

Env or VLP expression from A549 cells lines was induced by incubation in either 1 or 2 µg/ml doxycycline for 48 h. Then cell lysates, extracellular vesicles and VLPs were prepared as described above.

### Immunoprecipitation of Env from the surface of cells and VLPs

HEK 293T cells expressing Env, Rev and Tat or doxycycline-induced A549-Env cells were washed with 1x PBS. The cells were then incubated with 10 µg/ml anti-Env antibody or soluble CD4 (sCD4-Ig) for 1.5 h at room temperature. After three washes in 1x PBS, the cells were lysed (1x PBS/0.5% NP-40/protease inhibitors (Roche)) for 5 min on ice. The lysates were cleared by centrifugation at 13,200 x g for 10 min at 4°C, and the clarified supernatants were rotated during incubation with Protein A-Sepharose beads for 1 h at room temperature. The beads were pelleted (1,000 rpm for 1 min) and washed two times with wash buffer (1x PBS and 0.5% NP-40) and a third time with 1x PBS without NP-40. The beads were resuspended in 1x PBS/LDS/DTT and analyzed by Western blotting. To prepare the Input sample, an equal number of Env-expressing cells were lysed; cell lysates were cleared and analyzed by Western blotting, as above.

To evaluate Env antigenicity on the surface of VLPs, aliquots of virus particles (pelleted and resuspended in 1x PBS) were incubated with a panel of antibodies at 10 µg/mL concentration for 1.5 h at room temperature. One mL of chilled 1x PBS was then added and samples were centrifuged at 14,000 x g for 1 h at 4°C. The pellets were lysed in 100 µL chilled 1x PBS/0.5% NP-40/protease inhibitor cocktail. Lysates were rotated during incubation with Protein A-agarose beads for 1 h at 4°C and washed two times with chilled 1x PBS/0.1% NP-40 and a third time with 1x PBS without NP-40. The beads were resuspended in 1x PBS/LDS/DTT and used for Western blotting. To prepare the Input sample, an equal volume of virus suspension was mixed with 1 mL chilled 1x PBS and centrifuged at 14,000 x g for 1 h at 4°C; the pellet was resuspended in 1x PBS/LDS/DTT and analyzed by Western blotting.

### Deglycosylation of Env

Cells, VLPs and virions were lysed in 1x PBS/0.5% NP-40. The viral lysate was then boiled in denaturing buffer (New England BioLabs) for 10 min and treated with PNGase F or Endo Hf enzymes (New England BioLabs) for 1.5 h at 37°C according to the manufacturer’s protocol. The treated proteins were analyzed by reducing SDS-PAGE and Western blotting.

### Analysis of Env and SERINC5 by Western blotting

Clarified cell lysates, extracellular vesicles and pelleted virus or virus-like particles were analyzed by Western blotting using a nitrocellulose membrane and wet transfer (350 A, 75 min, Bio-Rad). Western blots were developed with 1:2,000 goat anti-gp120 polyclonal antibody (Invitrogen), 1:2,000 4E10 anti-gp41 antibody, 1:1,000 mouse anti-p24 monoclonal antibody (NIH HIV Reagent Program), 1:10,000 rabbit anti-hsp70 (K-20) antibody (Santa Cruz Biotechnology). The HRP-conjugated secondary antibodies were 1:2,000 rabbit anti-goat (Invitrogen), 1:2,000 goat anti-human (Invitrogen), and 1:10,000 goat anti-rabbit (Sigma-Aldrich). The intensity of protein bands on nonsaturated Western blots was quantified using ImageJ software.

To Western blot SERINC5, HEK293T cells were transfected with proviral constructs expressing the HIV-1_AD8_ and HIV-1_NL4-3_ Env variants along with plasmids expressing SERINC5/ΔSERINC5, or SERINC5 or ΔSERINC5 plasmids alone. The cell lysates were prepared as described above or the virions were pelleted and lysed. Western blots were developed using 1:250 rabbit polyclonal anti-SERINC5 antibody (Abcam), which detected an approximately 47-kDa SERINC5 band.

### Virus infectivity and cytopathic effect

HEK 293T cells were transfected with pNL4-3 plasmids containing infectious HIV-1 proviruses with the Envs of interest. Forty-eight hours after transfection, cell supernatants were collected and subjected to low-speed centrifugation (600 x g for 10 min at room temperature) to remove cell debris. Different volumes of the clarified cell supernatants were incubated with TZM-bl cells for 48 h in a 37°C/5% CO_2_ incubator after which luciferase activity in the cells was measured. Cell viability was measured using the CellTiter 96® Non-Radioactive Cell Proliferation Assay [(3-(4,5-dimethylthiazol-2-yl)-2,5-diphenyltetrazolium bromide (MTT)] kit from Promega. Briefly, different volumes of the clarified cell supernatants were incubated with TZM-bl cells for 48 h in a 37°C/5% CO_2_ incubator. Fifteen µL of dye solution (from the kit) was added to each well followed by incubation of the plate at 37°C for 4 h in a humidified CO_2_ incubator. One hundred µL of solubilization/stop solution (from the kit) was then added to each well. Absorbance at 570 nm was measured using a 96-well plate reader.

### Production of recombinant pseudoviruses expressing luciferase

As described previously (94,96), HEK 293T cells were transfected with pSVIIIenv plasmids expressing Env variants, the psPAX2 Gag-Pol packaging construct and the firefly luciferase-expressing HIV-1 vector at a 1:1:3 µg DNA ratio using effectene transfection reagent (Qiagen). Recombinant, luciferase-expressing viruses capable of a single round of replication were released into the cell medium and were harvested 48 h later. The virus-containing supernatants were clarified by low-speed centrifugation (600 x g for 10 min) and used for single-round infections.

To measure virus inhibition, the antibodies or the compounds (T20, BNM-III-170, BMS-806, amphotericin β-methyl ester and methyl-β-cyclodextrin) to be tested were incubated with pseudoviruses for 1 h at 37°C. The mixture was then added to Cf2Th-CD4/CCR5 target cells expressing CD4 and CCR5. Forty-eight hours later, the target cells were lysed, and the luciferase activity was measured. To test the effect of cholesterol oxidase on virus inhibition, the pseudoviruses were incubated with the enzyme for 5 h at 37°C and then the mixture was added to the target cells. To study the role of cholesterol on the mechanism of amphotericin β-methyl ester virus inhibition, the pseudotyped viruses were incubated with 1.5 mM or 3 mM MBCD or control buffer for 1 h at 37°C. The pseudotyped viruses were then pelleted, resuspended and incubated with amphotericin β-methyl ester at the indicated concentrations for 1 h at 37°C. The viruses were then added to Cf2Th-CD4/CCR5 cells. After two days of culture, the cells were lysed and luciferase activity was measured.

### Cell-cell fusion assay

For the alpha-complementation assay measuring cell-cell fusion, COS-1 effector cells were seeded in black-and-white 96-well plates and then cotransfected with plasmids expressing α-gal, Env variants and Tat at a 1:1:0.125 ratio, using Lipofectamine 3000 transfection reagent (Thermo Fisher Scientific) following the manufacturer’s protocol. At the same time, Cf2Th-CD4/CCR5 cells target cells in 6-well plates were cotransfected with a plasmid expressing ω-gal using Lipofectamine 3000 transfection reagent. Forty-eight hours after transfection, target cells were detached and resuspended in DMEM medium. Medium was aspirated from the effector cells, and target cell suspensions in 50-µL volumes were added to the effector cells (one target-cell well provides sufficient cells for 50 effector-cell wells). Plates were spun at 500 x g for 3 min and then incubated at 37°C in 5% CO_2_ for 2 h. Medium was removed, and cells were lysed in Tropix lysis buffer (Thermo Fisher Scientific). The β-galactosidase activity in the cell lysates was measured using the Galacto-Star Reaction Buffer Diluent with Galacto-Star Substrate (Thermo Fisher Scientific), according to the manufacturer’s instructions.

### Statistics

The concentrations of HIV-1 entry inhibitors that inhibit 50% of infection (IC_50_ values) were determined by fitting the data in five-parameter dose-response curves using GraphPad Prism 9. Unpaired t tests were performed and p values were calculated using GraphPad Prism 9. Correlation was evaluated by a Spearman rank correlation test.

### Data availability

The data reported herein are available from the corresponding author upon request. No new code or database depositions were associated with this research.

## Acknowledgments

We thank Ms. Elizabeth Carpelan for manuscript preparation. Antibodies against HIV-1 were kindly supplied by Dennis Burton (Scripps), Peter Kwong and John Mascola (Vaccine Research Center NIH), Barton Haynes (Duke University), Hermann Katinger (Polymun), James Robinson (Tulane University), and Marshall Posner (Mount Sinai Medical Center). The plasmid expressing SERINC5 was kindly supplied by Heinrich Göttlinger (UMass Worcester). The psPAX2 plasmid was a gift from Didier Trono (École Polytechnique Fédérale de Lausanne). We thank the NIH HIV Reagent program for providing reagents.

This study was supported by grants from the National Institutes of Health (grants AI145547, AI124982, AI176904, AI148379 and AI178833), by Michael Siff Funds for Basic Research (Dana-Farber Cancer Institute), by the Basic Science Core of the University of Alabama Center for AIDS Research (AI27767), and by a gift from the late William F. McCarty-Cooper.

We declare there are no conflicts of interest.

